# Proteomic indicators of oxidation and hydration state in colorectal cancer

**DOI:** 10.1101/035857

**Authors:** Jeffrey M. Dick

**Author notes:** Corresponding author: Jeffrey M. Dick.

## Abstract

New integrative approaches are needed to harness the potential of rapidly growing datasets of protein expression and microbial community composition in colorectal cancer (CRC). Chemical and thermodynamic models offer theoretical tools to describe populations of biomacromolecules and their relative potential for formation in different microenvironmental conditions. The average oxidation state of carbon (Z_C_) can be calculated as an elemental ratio from the chemical formulas of proteins, and water demand per residue 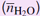 is computed by writing the overall formation reactions of proteins from basis species. Using reported results from clinical proteomic studies and microbial community profiling, many datasets exhibit higher mean Z_C_ or 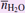 of proteins in carcinoma or adenoma compared to normal tissue. Microbial protein compositions have lower Z_C_ in bacteria enriched in fecal samples from cancer patients. In thermodynamic calculations, the potential for formation of the cancer-related proteins is energetically favored by changes in the chemical activity of H_2_O and fugacity of O_2_ that reflect the compositional differences. The compositional analysis suggests that a systematic shift in chemical composition is an essential feature of the cancer proteome, and the thermodynamic descriptions show that the observed proteomic transformations could be promoted by microenvironments shifted toward more oxidizing conditions and increased hydration levels.

## 1 INTRODUCTION

Datasets for differentially expressed proteins in cancer are often interpreted from a mechanistic perspective that emphasizes molecular interactions. Alternative approaches exemplified by recent models that use information theory demonstrate the possibility of interpreting proteomic expression data in a high-level conceptual framework (Rietman et al., 2016). These approaches may combine concepts from dynamical systems theory and thermodynamics, such as the correspondence of “attractor states” in landscape models with low-energy states of a system (Enver et al., 2009; Davies et al., 2011). Despite these advances, energetic functions for differential protein expression have not previously been formulated in terms of physicochemical variables that reflect the conditions of tumor microenvironments. The coupling of recent proteomic data with thermodynamic models using chemical components provides new perspectives on microenvironmental conditions that are conducive to carcinogenesis or healthy growth.

The purpose of the present study is to explore human proteomic and microbial community data for colorectal cancer within a chemical and thermodynamic framework using variables that represent changes in oxidation and hydration state. This is carried out first by comparing chemical compositions of up- and down-expressed proteins along the normal tissue-adenoma-carcinoma progression. Then, a thermodynamic model is used to quantify the overall energetics of the proteomic transformations in terms of chemical potential variables. This approach reveals not only common patterns of chemical changes among many proteomic datasets, but also the possibility that proteomic transformations may be shaped by energetic constraints associated with the changing tumor microenvironment.

Years of study of colorectal cancer (CRC), one of the most common types of human cancer, have resulted in the theory of genetic transformation as the primary driver of cancer progression (Kinzler and Vogelstein, 1996). However, not only multistep genetic changes, but also microenvironmental dynamics can influence cancer progression (Schedin and Elias, 2004). Many reactions in the microenvironment, such as those involving hormones or cell-cell signaling interactions, operate on fast timescales, but local hypoxia in tumors and other microenvironmental changes can develop and persist over longer timescales. Thus, the long timescales of carcinogenesis give cells sufficient time to adapt their proteomes to the differential energetic costs of biomolecular synthesis imposed by changing chemical conditions.

One of the characteristic features of tumors is varying degrees of hypoxia (Höckel and Vaupel, 2001). Hypoxic conditions promote activation of hypoxia-inducible genes by the HIF-1 transcription factor and intensify the mitochondrial generation of reactive oxygen species (ROS) (Murphy, 2009), leading to oxidative stress (Höckel and Vaupel, 2001; Semenza, 2008). It is important to note that there is significant intra-tumor and inter-tumor heterogeneity of oxygenation levels (Höckel and Vaupel, 2001; DeBerardinis and Cheng, 2010). Cancer cells can also exhibit changes in oxidation-reduction (redox) state; for example, redox potential (Eh) monitored *in vivo* in a fibrosarcoma cell line is altered compared to normal fibroblasts (Hutter et al., 1997).

The hydration states of cancer cells and tissues may also vary considerably from their healthy counterparts. Microwave detection of differences in dielectric constant resulting from greater water content in malignant tissue is being developed for medical imaging of breast cancer (Grzegorczyk et al., 2012). IR and Raman spectroscopic techniques also detect a greater hydration state of cancerous breast tissue, resulting from interaction of water molecules with hydrophilic cellular structures of cancer cells but negligible association with the triglycerides and other hydrophobic molecules that are more common in normal tissue (Abramczyk et al., 2014).

Increased hydration levels are associated with a higher abundance of hyaluronan found in the extracellular matrix (ECM) of migrating and metastatic cells (Toole, 2002). A higher subcellular hydration state may alter cell function by acting as a signal for protein synthesis and cell proliferation (Häussinger, 1996). It has been hypothesized that the increased hydration of cancer cells underlies a reversion to a more embryonic state (McIntyre, 2006). Based on all of these considerations, compositional and thermodynamic variables related to redox and hydration state have been selected as the primary descriptive variables in this study.

As noted by others, it is paradoxical that hypoxia, i.e. low oxygen partial pressure, could be a driving force for the generation of oxidative molecules. Possibly, the mitochondrial generation of ROS is a cellular mechanism for oxygen sensing (Guzy and Schumacker, 2006). Whether through hypoxia-induced oxidative stress or other mechanisms, proteins in cancer have been found to have a variety of oxidative post-translational modifications (PTM), including carbonylation and oxidation of cysteine residues (Yeh et al., 2010; Yang et al., 2013). Although proteome-level assessments of oxidative PTM are gaining traction (Yang et al., 2013), existing large-scale proteomic datasets may carry other signals of oxidation state. One possible “syn-translational” indicator of oxidation state, determined by the amino acid sequences of proteins, is the average oxidation state of carbon, introduced below. At the outset, it is not clear whether such a metric of oxidation state would more closely track hypoxia (i.e. relatively reducing conditions) that may arise in tumors, or a more oxidizing potential connected with ROS and oxidative PTM.

Density functional theory and other computational methods that yield electron density maps of proteins with known structure can be used to compute the partial charges, or oxidation states, of all the atoms. Spectroscopic methods can also be used to determine oxidation states of atoms in molecules (Gupta et al., 2014). These theoretical and empirical approaches offer the greatest precision in an oxidation state calculation, but it is difficult to apply them to the hundreds of proteins, many with undetermined three-dimensional structures, found to have significantly altered expression in proteomic experiments. Other methods for estimating the oxidation states of atoms in molecules may be needed to assess the overall direction of electron flow in a proteomic transformation.

Some textbooks of organic chemistry present the concept of formal oxidation states, in which the electron pair in a covalent bond is formally assigned to the more electronegative of the two atoms (e.g. Hendrickson et al., 1970, ch. 18). This rule is consistent with the IUPAC recommendations for calculating oxidation state of atoms in molecules, but generalizes the current IUPAC definitions such that the oxidation states of different carbon atoms in organic molecules can be distinguished (e.g. Loock, 2011; Gupta et al., 2014). In the primary structure of a protein, where no metal atoms are present and heteroatoms are bonded only to carbon, the average oxidation state of carbon (Z_C_) can be calculated as an elemental ratio, which is easily obtained from the amino acid composition (Dick, 2014). In a protein with the chemical formula C_*c*_H_*h*_N_*n*_O_*o*_S_*s*_, the average oxidation state of carbon (Z_C_) is

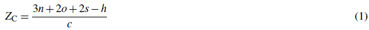

This equation is equivalent to others, also written in terms of numbers of the elements C, H, N, O and S, used for the average oxidation state of carbon in algal biomass (Bohutskyi et al., 2015), in humic and fulvic acids (Fekete et al., 2012), and for the nominal oxidation state of carbon in dissolved organic matter (Riedel et al., 2012).

Comparing the average carbon oxidation states in organic molecules is useful for quantifying the reactions of complex mixtures of organic matter in aerosols (Kroll et al., 2011), the growth of biomass (Hansen et al., 1994) and the production of biofuels (Borak et al., 2013; Bohutskyi et al., 2015). There is a considerable range of the average oxidation state of carbon in different amino acids (Masiello et al., 2008; Amend et al., 2013), with consequences for the energetics of synthesis depending on environmental conditions (Amend and Shock, 1998). Similarly, the nominal oxidation state of carbon can be used as a proxy for the standard Gibbs energies of oxidation reactions of various organic and biochemical molecules (Arndt et al., 2013). The oxidation state concept is easily applied as a bookkeeping tool to understand electron flow in metabolic pathways, yet may receive limited coverage in biochemistry courses (Halkides, 2000). There has been scant attention in the literature to the differences in carbon oxidation state among proteins or other biomacromolecules. Nevertheless, the ease of computation makes Z_C_ a useful metric for rapidly ascertaining the direction and magnitude of electron flow associated with proteomic transformations during disease progression.

Comparisons of oxidation states of carbon can be used to rank the energetics of reactions of organic molecules in particular systems (Amend et al., 2013). However, quantifying the energetics and mass-balance requirements of chemical transformations requires a more complete thermodynamic model. Thermodynamic models that are based on chemical components, i.e. a minimum number of independent chemical formula units that can be combined to form any chemical species in the system, have an established position in geochemistry (Anderson, 2005; Bethke, 2008). The implications of choosing different sets of components, called the “basis” (Bethke, 2008), have received relatively little discussion in biochemistry, although Alberty (2004) in this context highlighted the observation made by Callen (1985) that “[t]he choice of variables in terms of which a given system is formulated, while seemingly an innocuous step, is often the most crucial step in the solution”. Models built with different choices of components nevertheless yield equivalent results when consistently parameterized (Morel and Hering, 1993; Ravi Kanth et al., 2014). Accordingly, components are a type of chemical accounting for reactions in a system (Morel and Hering, 1993), and do not necessarily coincide with mechanistic models for those reactions.

The structure and dynamics of the hydration shells of proteins have important biological consequences (Levy and Onuchic, 2006) and can be investigated in molecular simulation studies (Wedberg et al., 2012). Statistical thermodynamics can be used to assess the effects of preferential hydration of protein surfaces on unfolding or other conformational changes (Lazaridis and Karplus, 2003). However, there is also a role for H_2_O as a chemical component in stoichiometric reactions representing the mass-balance requirements for formation of proteins with different amino acid sequences.

For example, a system of proteins composed of C, H, N, O and S can be described using the (non-innocuous) components CO_2_, NH_3_, H_2_S, O_2_ and H_2_O. Accordingly, stoichiometric reactions representing the formation of certain proteins at the expense of others during a proteomic transformation generally have non-zero coefficients on O_2_, H_2_O and the other components. These stoichiometric reactions can be written without specific knowledge of electron density or hydration by molecular H_2_O.

It bears repeating that reactions written using chemical components are not mechanistic representations. Instead, these reactions are specific statements of mass balance that are a requirement for thermodynamic models of chemically reacting systems (Helgeson et al., 2009). Flux-balance models of metabolic networks integrate stoichiometric constraints (e.g. Hiller and Metallo, 2013), but stoichiometric descriptions of proteomic transformations are less common, perhaps because of a greater degree of abstraction away from elementary reactions. Nevertheless, the differentially down- and up-expressed proteins in proteomic datasets can be viewed as representing the initial and final states of a chemically reacting system, which is then amenable to thermodynamic modeling.

The chemical potentials of components can be used to describe the internal state of a system and, for an open system, its relation to the environment. Oxygen fugacity is a variable that is related to the chemical potential of O_2_; it does not necessarily reflect the concentration of O_2_, but instead indicates the distribution of species with different oxidation states (Albarède, 2011). Theoretical calculation of the energetics of reactions as a function of oxygen fugacity provides a useful reference for the relative stabilities of organic molecules in different environments (Helgeson et al., 2009; Amend et al., 2013). However, in a cellular context a multidimensional approach may be required to quantify possible microenvironmental influences on the potentials for biochemical transformations. Likely variables include not only oxidation state but also water activity. Scenarios of early metabolic and cellular evolution (Pace, 1991; Russell and Hall, 1997; Damer and Deamer, 2015) lend additional support to the choice of water activity as a primary variable of interest.

A thermodynamic model that is formulated in terms of carefully selected components (basis species) affords a convenient description of a system. As described in the Methods, a basis is selected that reduces the empirical correlation between average oxidation state of carbon and the coefficient on H_2_O in formation reactions of proteins from basis species. The first part of the Results shows compositional comparisons for human and microbial proteins (Sections 3.1–3.2) in 35 datasets from 20 different studies. Many of the comparisons reveal higher mean Z_C_ or higher water demand for formation of proteins with higher expression in cancer compared to normal tissue. Contrary to the trend observed for human proteins, the mean protein compositions of bacteria enriched in cancer tend to have lower Z_C_.

To better understand the biochemical context of these differences, calculations reported in the second part of the Results use chemical affinity (negative Gibbs energy of reaction) to predict the most stable molecules as a function of oxygen fugacity and water activity (Sections 3.3–3.5). Mapping the theoretically calculated relative stabilities of proteins builds on the compositional descriptions toward quantifying the microenvironmental conditions that may promote or impede cancer progression.

## 2 METHODS

### 2.1 Data sources

This section describes the data sources and additional data processing steps applied in this study. An attempt was made to locate all currently available proteomic studies for clinical tissue on CRC including, among others, those listed in the “Tissue” and “Tissue subproteomes” sections of the review paper by de Wit et al. (2013) and in Supporting Table 3 (“Clinical Samples”) of the review paper by Martínez-Aguilar et al. (2013). To make the comparisons more robust, only datasets with at least 30 proteins in each of the up-and down-regulated groups were considered; however, all datasets from a given study were included if at least one of the datasets met this criterion. The reference keys for the selected studies shown below and in Table 1 are derived from the names of the authors and year of publication.

In comparisons between groups of up- and down-expressed proteins, the convention in this study is to consider proteins with higher expression in normal tissue or less-advanced cancer stages as a “normal” group (group 1), with number of proteins *n*_1_, while proteins with higher expression in cancer or more-advanced cancer stages are categorized as a “cancer” group (group 2), with number of proteins *n*_2_. Accordingly, in the dataset of Uzozie et al. (2014) comparing normal mucosa and adenoma, the proteins up-expressed in adenoma are assigned to group 2, while in the adenoma-carcinoma dataset of Knol et al. (2014), the proteins with higher expression in adenoma are assigned to group 1 (see Table 1).

Names or IDs of genes or proteins given in the sources were searched in UniProt. The corresponding UniProt IDs are provided in the `*.csv` data files in Dataset S1. Amino acid sequences of human proteins were taken from the UniProt reference proteome (files `UP000005640_9606.fasta.gz` containing canonical, manually reviewed sequences, and `UP000005640_9606_additional.fasta.gz` containing isoforms and unreviewed sequences, dated 2016-04-13, downloaded from ftp://ftp.uniprot.org/pub/databases/uniprot/current_release/knowledgebase/reference_proteomes/Eukaryota/). Entire sequences were used; i.e., signal and propeptides were not removed when calculating the amino acid compositions. However, amino acid compositions were calculated for particular isoforms, if these were identified in the sources. Files `human.aa.csv` and `human_additional.aa.csv` in Dataset S1 contain the amino acid compositions of the proteins calculated from the UniProt reference proteome. In a few cases, amino acid compositions of unreviewed or obsolete sequences in UniProt, not available in the reference proteome, were individually compiled; these are contained in file `human2.aa.csv` in Dataset S1.

Reported gene names were converted to UniProt IDs using the UniProt mapping tool http://www.uniprot.org/mapping, and IPI accession numbers were converted to UniProt IDs using the DAVID conversion tool http://david.ncifcrf.gov/content.jsp?file=conversion.html. For proteins with no automatically generated matches, manual searches in UniProt of the protein descriptions, where available, were performed. Proteins with missing or duplicated identifiers, or those that could not be matched to a UniProt ID, were omitted from the comparisons here. Therefore, the numbers of proteins actually used in the comparisons (listed in Table 1) may be different from the numbers of proteins reported by the authors and summarized below.

`WTK+08`: Watanabe et al. (2008) used 2-nitrobenzenesulfenyl labeling and MS/MS analysis to identify 128 proteins with differential expression in paired CRC and normal tissue specimens from 12 patients. The list of proteins used in this study was generated by combining the lists of up- and down-regulated proteins from Table 1 and Supplementary Data 1 of Watanabe et al. (2008) with the Swiss-Prot and UniProt accession numbers from their Supplementary Data 2.

`AKP+10`: Albrethsen et al. (2010) used nano-LC-MS/MS to characterize proteins from the nuclear matrix fraction in samples from 2 patients each with adenoma (ADE), chromosomal instability CRC (CIN+) and microsatellite instability CRC (MIN+). Cluster analysis was used to classify proteins with differential expression between ADE and CIN+, MIN+, or in both subtypes of carcinoma (CRC). Here, gene names from Supplementary Table 5-7 of Albrethsen et al. (2010) were converted to UniProt IDs using the UniProt mapping tool.

`JKMF10`: Jimenez et al. (2010) compiled a list of candidate serum biomarkers from a meta-analysis of the literature. In the meta-analysis, 99 up- or down-expressed proteins were identified in at least 2 studies. The list of UniProt IDs used in this study was taken from Table 4 of Jimenez et al. (2010).

`XZC+10`: Xie et al. (2010) used a gel-enhanced LC-MS method to analyze proteins in pooled tissue samples from 13 stage I and 24 stage II CRC patients and pooled normal colonic tissues from the same patients. Here, IPI accession numbers from Supplemental Table 4 of Xie et al. (2010) were converted to UniProt IDs using the DAVID conversion tool.

`ZYS+10`: Zhang et al. (2010) used acetylation stable isotope labeling and LTQ-FT MS to analyze proteins in pooled microdissected epithelial samples of tumor and normal mucosa from 20 patients, finding 67 and 70 proteins with increased and decreased expression (ratios ≥20 or ≤0.5). Here, IPI accession numbers from Supplemental Table 4 of Zhang et al. (2010) were converted to UniProt IDs using the DAVID conversion tool.

`BPV+11`: Besson et al. (2011) analyzed microdissected cancer and normal tissues from 28 patients (4 adenoma samples and 24 CRC samples at different stages) using iTRAQ labeling and MALDI-TOF/TOF MS to identify 555 proteins with differential expression between adenoma and stage I, II, III, IV CRC. Here, gene names from supplemental Table 9 of Besson et al. (2011) were converted to UniProt IDs using the UniProt mapping tool.

`JCF+11`: Jankova et al. (2011) analyzed paired samples from 16 patients using iTRAQ-MS to identify 118 proteins with >1.3-fold differential expression between CRC tumors and adjacent normal mucosa. The protein list used in this study was taken from Supplementary Table 2 of Jankova et al. (2011).

`MRK+11`: Mikula et al. (2011) used iTRAQ labeling with LC-MS/MS to identify a total of 1061 proteins with differential expression (fold change ≥1.5 and false discovery rate ≤0.01) between pooled samples of 4 normal colon (NC), 12 tubular or tubulo-villous adenoma (AD) and 5 adenocarcinoma (AC) tissues. The list of proteins used in this study was taken from from Table S8 of Mikula et al. (2011).

`KKL+12`: Kim et al. (2012) used difference in-gel electrophoresis (DIGE) and cleavable isotope-coded affinity tag (cICAT) labeling followed by mass spectrometry to identify 175 proteins with more than 2-fold abundance ratios between microdissected and pooled tumor tissues from stage-IV CRC patients with good outcomes (survived more than five years; 3 patients) and poor outcomes (died within 25 months; 3 patients). The protein list used in this study was made by filtering the cICAT data from Supplementary Table 5 of Kim et al. (2012) with an abundance ratio cutoff of >2 or <0.5, giving 147 proteins. IPI accession numbers were converted to UniProt IDs using the DAVID conversion tool.

`KYK+12`: Kang et al. (2012) used mTRAQ and cICAT analysis of pooled microsatellite stable (MSS-type) CRC tissues and pooled matched normal tissues from 3 patients to identify 1009 and 478 proteins in cancer tissue with increased and decreased expression by higher than 2-fold, respectively. Here, the list of proteins from Supplementary Table 4 of Kang et al. (2012) was filtered to include proteins with expression ratio >2 or <0.5 in both mTRAQ and cICAT analyses, leaving 175 up-expressed and 248 down-expressed proteins in CRC. Gene names were converted to UniProt IDs using the UniProt mapping tool.

`WOD+12`: Wisniewski et al. (2012) used LC-MS/MS to analyze proteins in microdissected samples of formalin-fixed paraffin-embedded (FFPE) tissue from 8 patients; at *P* < 0.01, 762 proteins had differential expression between normal mucosa and primary tumors. The list of proteins used in this study was taken from Supplementary Table 4 of Wiśniewski et al. (2012).

`YLZ+12`: Yao et al. (2012) analyzed the conditioned media of paired stage I or IIA CRC and normal tissues from 9 patients using lectin affinity capture for glycoprotein (secreted protein) enrichment by nano LC-MS/MS to identify 68 up-regulated and 55 down-regulated differentially expressed proteins. IPI accession numbers listed in Supplementary Table 2 of Yao et al. (2012) were converted to UniProt IDs using the DAVID conversion tool.

`MCZ+13`: Mu et al. (2013) used laser capture microdissection (LCM) to separate stromal cells from 8 colon adenocarcinoma and 8 non-neoplastic tissue samples, which were pooled and analyzed by iTRAQ to identify 70 differentially expressed proteins. Here, gi numbers listed in Table 1 of Mu et al. (2013) were converted to UniProt IDs using the UniProt mapping tool; FASTA sequences of 31 proteins not found in UniProt were downloaded from NCBI and amino acid compositions were added to `human2.aa.csv`.

`KWA+14`: Knol et al. (2014) used differential biochemical extraction to isolate the chromatin-binding fraction in frozen samples of colon adenomas (3 patients) and carcinomas (5 patients), and LC-MS/MS was used for protein identification and label-free quantification. The results were combined with a database search to generate a list of 106 proteins with nuclear annotations and at least a three-fold expression difference. Here, gene names from Table 2 of Knol et al. (2014) were converted to UniProt IDs.

`UNS+14`: Uzozie et al. (2014) analyzed 30 samples of colorectal adenomas and paired normal mucosa using iTRAQ labeling, OFFGEL electrophoresis and LC-MS/MS. 111 proteins with expression fold changes (log_2_) at least +/− 0.5 and statistical significance threshold *q* < 0.02 that were also quantified in cell-line experiments were classified as “epithelial cell signature proteins”. UniProt IDs were taken from Table III of Uzozie et al. (2014).

`WKP+14`: de Wit et al. (2014) analyzed the secretome of paired CRC and normal tissue from 4 patients, adopting a five-fold enrichment cutoff for identification of candidate biomarkers. Here, the list of proteins from Supplementary Table 1 of de Wit et al. (2014) was filtered to include those with at least five-fold greater or lower abundance in CRC samples and *p* < 0.05. Two proteins listed as “Unmapped by Ingenuity” were removed, and gene names were converted to UniProt IDs using the UniProt mapping tool.

`STK+15`: Sethi et al. (2015) analyzed the membrane-enriched proteome from tumor and adjacent normal tissues from 8 patients using label-free nano-LC-MS/MS to identify 184 proteins with a fold change > 1.5 and *p*-value < 0.05. Here, protein identifiers from Supporting Table 2 of Sethi et al. (2015) were used to find the corresponding UniProt IDs.

`WDO+15`: Wisniewski et al. (2015) analyzed 8 matched formalin-fixed and paraffin-embedded (FFPE) samples of normal tissue (N) and adenocarcinoma (C) and 16 nonmatched adenoma samples (A) using LC-MS to identify 2300 (N/A), 1780 (A/C) and 2161 (N/C) up- or down-regulated proteins at *p* < 0.05. The list of proteins used in this study includes only those marked as having a significant change in SI Table 3 of Wiśniewski et al. (2015).

`LPL+16`: Li et al. (2016) used iTRAQ and 2D LC-MS/MS to analyze pooled samples of stroma purified by laser capture microdissection (LCM) from 5 cases of non-neoplastic colonic mucosa (NC), 8 of adenomatous colon polyps (AD), 5 of colon carcinoma in situ (CIS) and 9 of invasive colonic carcinoma (ICC). A total of 222 differentially expressed proteins between NNCM and other stages were identified. Here, gene symbols from Supplementary Table S3 of Li et al. (2016) were converted to UniProt IDs using the UniProt mapping tool.

`PHL+16`: Peng et al. (2016) used iTRAQ 2D LC-MS/MS to analyze pooled samples from 5 cases of normal colonic mucosa (NC), 8 of adenoma (AD), 5 of carcinoma in *situ* (CIS) and 9 of invasive colorectal cancer (ICC). A total of 326 proteins with differential expression between two successive stages (and, for CIS and ICC, also differentially expressed with respect to NC) were detected. The list of proteins used in this study was generated by converting the gene names in Supplementary Table 4 of Peng et al. (2016) to UniProt IDs using the UniProt mapping tool.

### 2.2 Basis I

To formulate a thermodynamic description of a chemically reacting system, an important choice must be made regarding the *basis species* used to describe the system. The basis species, like thermodynamic components, are a minimum number of chemical formula units that can be linearly combined to generate the composition of any chemical species in the system of interest. Stated differently, any species can be formed by combining the components, but components can not be used to form other components (VanBriesen and Rittmann, 1999). Within these constraints, any specific choice of a basis is theoretically permissible. In making the choice of components, convenience (Gibbs, 1875), ease of interpretation and relationship with measurable variables, as well as availability of thermodynamic data (e.g. Helgeson, 1970), and kinetic favorability (May et al., 2001) are other useful considerations. Once the basis species are chosen, the stoichiometric coefficients in the formation reaction for any chemical species are algebraically determined.

Following previous studies (e.g. Dick, 2008), the basis species initially chosen here are CO_2_, H_2_O, NH_3_, H_2_S and O_2_ (Basis I). The reaction representing the overall formation from these basis species of a protein having the formula C_*c*_H_*h*_N_*n*_O_*o*_S_*s*_ is

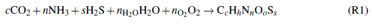

where 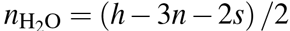 and 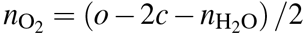. Dividing 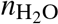 by the length of the protein gives the water demand per residue 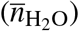, which is used here because proteins in the comparisons generally have different sequence lengths.

**Figure. 1.**
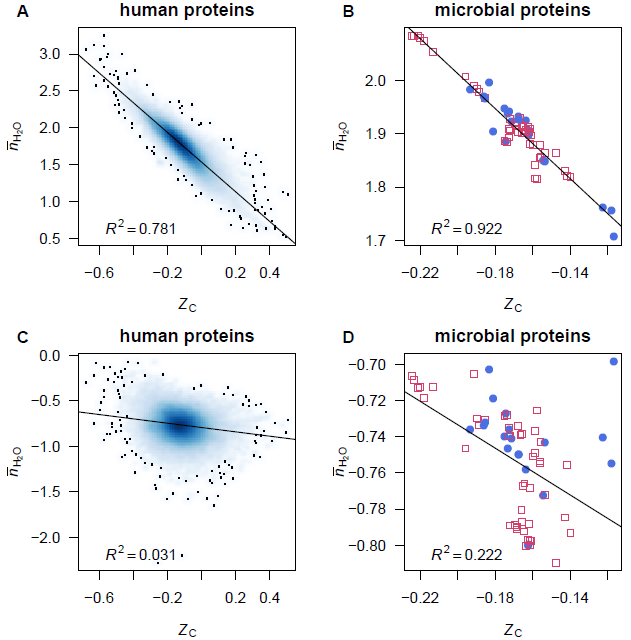
Scatterplots of average oxidation state of carbon (Z_C_) and water demand per residue (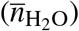). Data are plotted for (A,C) individual human proteins and (B,D) mean composition of proteins from microbial genomes, with 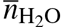 computed using (A,B) Basis I (Reaction R1) or (C,D) Basis II (Reaction R2). Linear least-squares fits and *R^2^* values are shown. In (A) and (C), the intensity of shading corresponds to density of points, produced using the `smoothScatter()` function of R graphics (R Core Team, 2016). The label in plot (A) identifies a particular protein, MUC1, which is used for the example calculations (see Reactions R3 and R4).

These or similar sets of inorganic species (such as H_2_ instead of O_2_) are often used in studying reaction energetics in geobiochemistry (e.g. Shock and Canovas, 2010). However, as seen in Fig. 1A and B, there is a high correlation between Z_C_ of protein molecules and 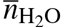 in the reactions to form the proteins from Basis I (note that the choice of basis species here affects only 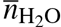 and not Z_C_, which is derived from an elemental ratio). Because of this stoichiometric *interdependence*, changing either redox or hydration potential, while holding the chemical potentials of the remaining basis species constant, have correlated effects on the energetics of chemical transformations (see Section 3.6 below). A different set of basis species can be chosen that reduces this correlation and is more useful for convenient description of subcellular processes.

### 2.3 Basis II

In this exploratory study, we restrict attention to at most two variables, with the implication that the others are held constant. In a subcellular setting, assuming that chemical potentials of CO_2_, NH_3_ and H_2_S do not change throughout a proteomic transformation, as implied by varying the chemical potentials of O_2_ and H_2_O in Basis I, may be less appropriate than assuming constant (or possibly buffered) potentials of more complex metabolites. In thermodynamic models for systems of proteins, constant chemical activities of chemical components having the compositions of amino acids might be a reasonable provision.

Although 1140 3-way combinations can be made of the 20 common proteinogenic amino acids, only 324 of the combinations contain cysteine and/or methionine (one of these is required to provide sulfur), and of these only 300, when combined with O_2_ and H_2_O, are compositionally independent. The slope, intercept and *R^2^* of the linear least-squares fits between Z_C_ and 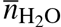 using each possible basis are listed in file `AAbasis.csv` in Dataset S1. Many of these combinations have lower *R*^2^ and lower slopes than found for Basis I (Fig. 1A, B), indicating a decreased correlation. From those with a lower correlation, but not the lowest, the basis including cysteine (Cys), glutamic acid (Glu), glutamine (Gln), O_2_ and H_2_O (Basis II) has been selected for use in this study. The scatter plots and fits between Z_C_ and 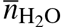 using Basis II are shown in Fig. 1C and D.

A secondary consideration in choosing this basis instead of others with even lower *R*^2^ is the centrality of glutamine and glutamic acid in many metabolic pathways (e.g. DeBerardinis and Cheng, 2010). Accordingly, these amino acids may be kinetically more reactive than others in pathways of protein synthesis and degradation. The presence of side chains derived from cysteine and glutamic acid in the abundant glutathione molecule (GSH), associated with redox homeostasis, is also suggestive of a central metabolic requirement for these amino acids. Again, it must be stressed that the current provisional choice of basis species is neither uniquely determined nor necessarily optimal. More experience with thermodynamic modeling and better biochemical intuition will likely provide reasons to refine these calculations using a different basis, perhaps including metabolites other than amino acids.

A general formation reaction using Basis II is

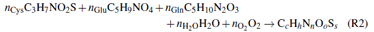

where the reaction coefficients (*n*_Cys_, *n*_Glu_, *n*_Gln_, 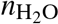 and 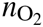) can be obtained by solving

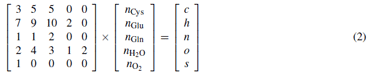

Although the definition of basis species requires that they are themselves compositionally non-degenerate, the matrix equation emphasizes the interdependence of the stoichiometric reaction coefficients. A consequence of this multiple dependence is that single variables such as 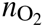 and 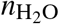 are not *simple* variables, but are influenced by both the intrinsic chemical makeup of the protein and the choice of basis species used to describe the system.

The combination of molecules shown in Reaction R2 does not represent the actual mechanism of synthesis of the proteins. Instead, reactions such as this allow for accounting of mass-conservation requirements and subsequent generation of a thermodynamic description of the effects of changing the local environment (i.e. chemical potentials of O_2_ and H_2_O) on the potential for formation of different proteins.

As an example of a specific calculation, consider the following reaction:

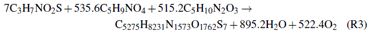

This reaction represents the overall formation from the basis species of one mole of the protein MUC1. This is a chromatin-binding protein that is highly up-expressed in CRC cells (Knol et al., 2014). The average oxidation state of carbon (Z_C_; Equation 1) in MUC1 is 0.005. Water is released in Reaction R3, so the *water demand* 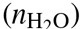 is negative. The length of this protein is 1255 amino acid residues, giving the water demand per residue, 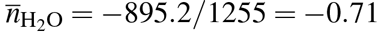. The value of Z_C_ indicates that MUC1 is a relatively highly oxidized protein, while its 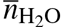 places it near the median water demand for cancer-associated proteins in this dataset (see Fig. 2 below).

### 2.4 Thermodynamic calculations

Standard molal thermodynamic properties of the amino acids and unfolded proteins estimated using amino acid group additivity were calculated as described by Dick et al. (2006), taking account of updated values for the methionine sidechain group (LaRowe and Dick, 2012). All calculations were carried out at 37 °C and 1 bar. The temperature dependence of standard Gibbs energies was calculated using the revised Helgeson-Kirkham-Flowers (HKF) equations of state (Helgeson et al., 1981; Tanger and Helgeson, 1988). Thermodynamic properties for O_2_ (gas) were calculated using data from Wagman et al. (1982) and the Maier-Kelley heat capacity function (Kelley, 1960). Properties of H_2_O (liquid) were calculated using data and extrapolations coded in Fortran subroutines from the SUPCRT92 package (Johnson et al., 1992), as provided in the CHNOSZ package (Dick, 2008).

Chemical affinities of reactions were calculated using activities of amino acids in the basis equal to 10 ^-4^, and activities of proteins equal to 1/(protein length) (i.e., unit activity of amino acid residues). The chemical affinities of formation of proteins are also sensitive to the environmental conditions represented by temperature (*T*), pressure (*P*) and the chemical potentials of basis species. Continuing with the example of Reaction R3, an estimate of the standard Gibbs energy 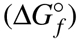 of the aqueous protein molecule (Dick et al., 2006; LaRowe and Dick, 2012) at 37 °C is −40974 kcal/mol; combined with the standard Gibbs energies of the basis species, this give a standard Gibbs energy of reaction 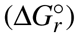 equal to 66889 kcal/mol. At 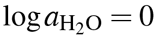 and 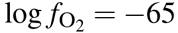, with activities of the amino acid basis species equal to 10^−4^, the overall Gibbs energy (Δ*G_r_*) is 24701 kcal/mol. The negative of this value is the chemical affinity (*A*) of the reaction. The per-residue chemical affinity (used in order to compare the relative stabilities of proteins of different sizes) for formation of protein MUC1 in the stated conditions is −19.7 kcal/mol. (This calculation can be reproduced using the function `reaction()` in file `plot.R` in Dataset S1.)

In a given system, proteins with higher (more positive) chemical affinity are relatively energetically stabilized, and theoretically have a higher propensity to be formed. Therefore, the differences in affinities reflect not only the amino acid compositions of the protein molecules but also the potential for local environmental conditions to influence the relative abundances of proteins.

### 2.5 Weighted rank difference

The contours on relative stability diagrams for the “normal” and “cancer” groups (see Fig. 6 below) depict the weighted rank differences of chemical affinities of the groups of proteins. To illustrate this calculation, consider a hypothetical system composed of 3 cancer (C) and 4 healthy (H) proteins. Suppose that under one set of conditions (i.e. specified 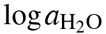 and 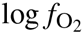), the per-residue affinities of the proteins give the following ranking in ascending order (I):

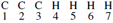

This gives as the sum of ranks for cancer proteins Σ *r*_C_ = 6, and for healthy proteins Σ *r*_H_ = 22. The difference in sum of ranks is Δ*r*_C−H_ = −16; the negative value is associated with a higher rank sum for the healthy proteins, indicating that these as a group are more stable than the cancer proteins. In a second set of conditions, we might have (II):

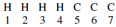

Here, the difference of rank sums is Δ*r*_C−H_ = 18 − 10 = 8.

For systems where the numbers of proteins in the two groups are equal, the maximum possible differences in rank sums would have equal absolute values, but that is not the case in this and other systems having unequal numbers of up- and down-expressed proteins. To characterize these datasets, the weighted rank-sumdifference can be calculated using

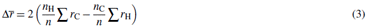

where *n*_H_, *n*_C_ and *n* are the numbers of healthy, cancer, and total proteins in the comparison. In the example here, we have *n*_H_/*n* = 4/7 and *n*_C_/*n* = 3/7. Eq. (3) then gives 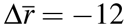 and 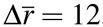, respectively, for conditions (I) and (II) above, showing equal weighted rank-sum differences for the two extreme rankings.

We can also consider a situation where the ranks of the proteins are evenly distributed:

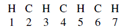

Here the absolute difference of rank sums is Δ*r*_C−H_; = 12 − 16 = −4, but the weighted rank-sum difference is 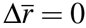. The zero value for an even distribution and the opposite values for the two extremes demonstrate the applicability of this weighting scheme.

### 2.6 Software availability

All statistical and thermodynamic calculations were performed using R (R Core Team, 2016). Thermodynamic calculations were carried out using R package CHNOSZ (Dick, 2008). Effect sizes (see below) were calculated using R package orddom (Rogmann, 2013). Figures were generated using CHNOSZ and graphical functions available in R together with the R package colorspace (Ihaka et al., 2015) for constructing an HCL-based color palette (Zeileis et al., 2009). With the mentioned packages installed, the figures in this paper can be reproduced using the code (`plot.R`) and data files (`*.csv`) in Dataset S1.

## 3 RESULTS

### 3.1 Compositional descriptions of human proteins

Comparisons of proteome composition in terms of average oxidation state of carbon (Z_C_) and water demand per residue 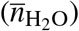 are presented in Fig. 2 and Table 1. Fig. 2 shows scatterplots of individual protein composition for proteomes in three representative studies. Each of these exhibits a strongly differential trend in Z_C_ or 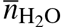 that can be visually identified. In Fig. 2A, chromatin-binding proteins highly expressed in carcinoma (Knol et al., 2014) as a group exhibit a lower Z_C_ than those found to be more abundant in adenoma. In Fig. 2B, proteins relatively highly expressed in epithelial cells in adenoma (Uzozie et al., 2014) tend to have higher 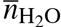 than the proteins more highly expressed in paired normal tissues. Differentially expressed proteins between adenoma and normal tissue identified in a recent deep-proteome analysis (Wisniewski et al., 2015) are compared in Fig. 2C, showing that proteins up-expressed in adenoma are relatively oxidized (i.e. have higher Z_C_).

In order to quantify these differences, Table 1 shows the numbers of proteins in each comparison (*n*_1_ for normal or less advanced cancer stage; *n*_2_ for tumor or more advanced cancer stage), differences of means (MD), common language effect size as percentages (ES), and *p*-values calculated using the Mann-Whitney-Wilcoxon test. This non-parametric test is suitable for data which may not be normally distributed. For a given experiment, the common language effect size, or probability of superiority, describes the probability that Z_C_ or 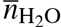 of a protein is higher in the cancer group than in the normal group. That is, percent values of the ES greater than (or less than) 50 indicate a greater proportion of pairwise higher (or lower) Z_C_ or 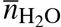 of proteins in the *n*_2_ compared to *n*_1_ groups. The CLES and *p*-value are used here to allow for a subjective assessment of the compositional differences. Arbitrarily, CLES values ≥60 or ≤40 and *p*-values < 0.05 are highlighted in the table. The corresponding mean differences are underlined for *p* < 0.05, or bolded if CLES is also ≥60 or ≤40. These arbitrary cutoffs highlight datasets with the largest and most significant differences in Z_C_ and 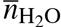. Mean and median values of Z_C_ and 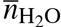 are given in file `summary.csv` in Dataset S1.

**Fig. 2.**
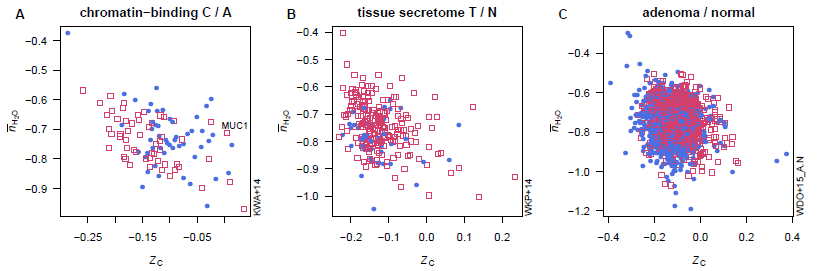
Average oxidation state of carbon (Z_C_) and water demand per residue 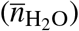 for proteins in selected datasets. Open red squares represent proteins enriched in tumors or more advanced cancer stages, and filled blue circles represent proteins enriched in normal tissue or less advanced cancer stages.

Counting the underlined and highlighted MD values in Table 1, the number of datasets with a significant difference in Z_C_ (18) is greater than those with a significant difference in 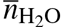(10). Of the 13 unique studies yielding at least one dataset with a significant difference in Z_C_, 8 exhibit a higher mean value in adenoma and/or carcinoma compared to normal tissue. Datasets from a couple of studies (Besson et al., 2011; Wisniewski et al., 2015) exhibit mean values of Z_C_ with opposite signs of the differences between adenoma or carcinoma compared normal tissue.

Most of the studies analyzed proteins in whole or microdissected tissue, but datasets from two other studies (both from the same laboratory) represent the nuclear matrix or chromatin-binding fraction (Albrethsen et al., 2010; Knol et al., 2014). These two datasets give lower mean Z_C_ of proteins more highly expressed in carcinoma than adenoma. Two other datasets have a lower mean value of Z_C_ in carcinoma (Albrethsen et al., 2010; Wiśniewski et al., 2015), and one has a higher mean value (Mikula et al., 2011).

**Table I.**
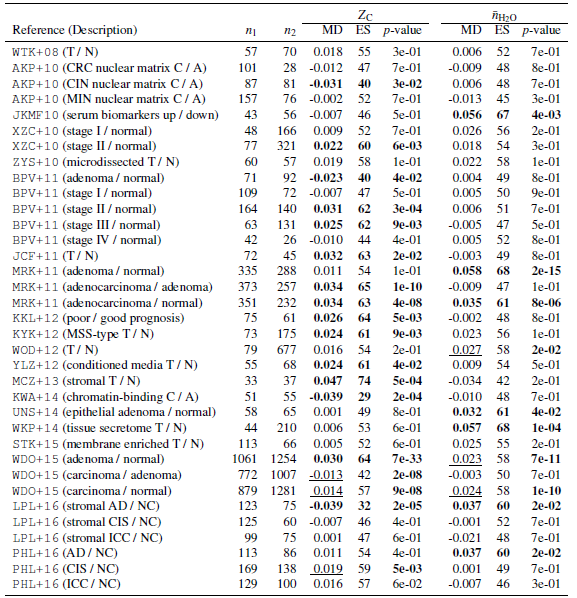
Summary of compositional comparisons for human proteins. Mean differences (MD), percent values of common language effect size (ES), and *p*-values are shown for comparisons between groups of *n*_1_ and *n*_2_ proteins reported to have higher abundance in normal and cancer tissue (or less and more advanced cancer stage), respectively. The textual descriptions are written such that the ordering around the slash (“/”) corresponds to *n*_2_ / *n*_1_ Abbreviations: T / N (tumor / normal), C / A (carcinoma / adenoma). References and specific abbreviations used in the descriptions are given in Section 2.1.

| Reference (Description) | $n_1$ | $n_2$ | $Z_C$ | | | $\bar{n}_{H_2O}$ | | |
| --- | --- | --- | --- | --- | --- | --- | --- | --- |
| | | | MD | ES | $p$-value | MD | ES | $p$-value |
| WTK+08 (T / N) | 57 | 70 | 0.018 | 55 | 3e-01 | 0.006 | 52 | 7e-01 |
| AKP+10 (CRC nuclear matrix C / A) | 101 | 28 | -0.012 | 47 | 7e-01 | -0.009 | 48 | 8e-01 |
| AKP+10 (CIN nuclear matrix C / A) | 87 | 81 | **-0.031** | **40** | **3e-02** | 0.006 | 48 | 7e-01 |
| AKP+10 (MIN nuclear matrix C / A) | 157 | 76 | -0.002 | 52 | 7e-01 | -0.013 | 45 | 3e-01 |
| JKMF10 (serum biomarkers up / down) | 43 | 56 | -0.007 | 46 | 5e-01 | **0.056** | **67** | **4e-03** |
| XZC+10 (stage I / normal) | 48 | 166 | 0.009 | 52 | 7e-01 | 0.026 | 56 | 2e-01 |
| XZC+10 (stage II / normal) | 77 | 321 | **0.022** | **60** | **6e-03** | 0.018 | 54 | 3e-01 |
| ZYS+10 (microdissected T / N) | 60 | 57 | 0.019 | 58 | 1e-01 | 0.022 | 58 | 1e-01 |
| BPV+11 (adenoma / normal) | 71 | 92 | **-0.023** | **40** | **4e-02** | 0.004 | 49 | 8e-01 |
| BPV+11 (stage I / normal) | 109 | 72 | -0.007 | 47 | 5e-01 | 0.005 | 50 | 9e-01 |
| BPV+11 (stage II / normal) | 164 | 140 | **0.031** | **62** | **3e-04** | 0.006 | 51 | 7e-01 |
| BPV+11 (stage III / normal) | 63 | 131 | **0.025** | **62** | **9e-03** | -0.005 | 47 | 5e-01 |
| BPV+11 (stage IV / normal) | 42 | 26 | -0.010 | 44 | 4e-01 | 0.005 | 52 | 8e-01 |
| JCF+11 (T / N) | 72 | 45 | **0.032** | **63** | **2e-02** | -0.003 | 49 | 8e-01 |
| MRK+11 (adenoma / normal) | 335 | 288 | 0.011 | 54 | 1e-01 | **0.058** | **68** | **2e-15** |
| MRK+11 (adenocarcinoma / adenoma) | 373 | 257 | **0.034** | **65** | **1e-10** | -0.009 | 47 | 1e-01 |
| MRK+11 (adenocarcinoma / normal) | 351 | 232 | **0.034** | **63** | **4e-08** | **0.035** | **61** | **8e-06** |
| KKL+12 (poor / good prognosis) | 75 | 61 | **0.026** | **64** | **5e-03** | -0.002 | 48 | 8e-01 |
| KYK+12 (MSS-type T / N) | 73 | 175 | **0.024** | **61** | **9e-03** | 0.023 | 56 | 1e-01 |
| WOD+12 (T / N) | 79 | 677 | 0.016 | 54 | 2e-01 | <u>0.027</u> | 58 | **2e-02** |
| YLZ+12 (conditioned media T / N) | 55 | 68 | **0.024** | **61** | **4e-02** | 0.009 | 54 | 5e-01 |
| MCZ+13 (stromal T / N) | 33 | 37 | **0.047** | **74** | **5e-04** | -0.034 | 42 | 2e-01 |
| KWA+14 (chromatin-binding C / A) | 51 | 55 | **-0.039** | **29** | **2e-04** | -0.010 | 48 | 7e-01 |
| UNS+14 (epithelial adenoma / normal) | 58 | 65 | 0.001 | 49 | 8e-01 | **0.032** | **61** | **4e-02** |
| WKP+14 (tissue secretome T / N) | 44 | 210 | 0.006 | 53 | 6e-01 | **0.057** | **68** | **1e-04** |
| STK+15 (membrane enriched T / N) | 113 | 66 | 0.005 | 52 | 6e-01 | 0.025 | 55 | 2e-01 |
| WDO+15 (adenoma / normal) | 1061 | 1254 | **0.030** | **64** | **7e-33** | <u>0.023</u> | 58 | **7e-11** |
| WDO+15 (carcinoma / adenoma) | 772 | 1007 | <u>-0.013</u> | 42 | **2e-08** | -0.003 | 50 | 7e-01 |
| WDO+15 (carcinoma / normal) | 879 | 1281 | <u>0.014</u> | 57 | **9e-08** | <u>0.024</u> | 58 | **1e-10** |
| LPL+16 (stromal AD / NC) | 123 | 75 | **-0.039** | **32** | **2e-05** | **0.037** | **60** | **2e-02** |
| LPL+16 (stromal CIS / NC) | 125 | 60 | -0.007 | 46 | 4e-01 | -0.001 | 52 | 7e-01 |
| LPL+16 (stromal ICC / NC) | 99 | 75 | 0.001 | 47 | 6e-01 | -0.021 | 48 | 7e-01 |
| PHL+16 (AD / NC) | 113 | 86 | 0.011 | 54 | 4e-01 | **0.037** | **60** | **2e-02** |
| PHL+16 (CIS / NC) | 169 | 138 | <u>0.019</u> | 59 | **5e-03** | 0.001 | 49 | 7e-01 |
| PHL+16 (ICC / NC) | 129 | 100 | 0.016 | 57 | 6e-02 | -0.007 | 46 | 3e-01 |

**Table II.**
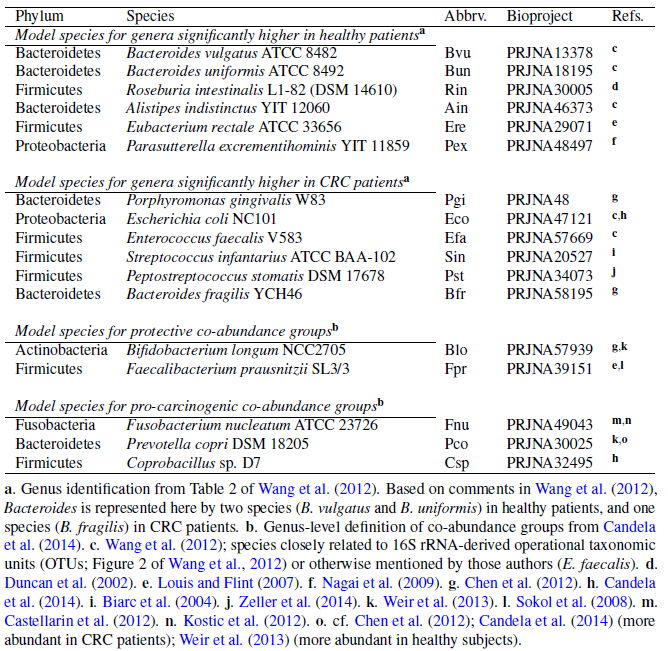
Microbial species selected as models for genera and co-abundance groups that differ between CRC and healthy patients. **a**. Genus identification from Table 2 of Wang et al. (2012). Based on comments in Wang et al. (2012), Bacteroides is represented here by two species (*B. vulgatus* and *B. uniformis*) in healthy patients, and one species (*B. fragilis*) in CRC patients. **b**. Genus-level definition of co-abundance groups from Candela et al. (2014). **c**. Wang et al. (2012); species closely related to 16S rRNA-derived operational taxonomic units (OTUs; Figure 2 of Wang et al., 2012) or otherwise mentioned by those authors (*E. faecalis*). **d**. Duncan et al. (2002). **e**. Louis and Flint (2007). **f**. Nagai et al. (2009). **g**. Chen et al. (2012). **h**. Candela et al. (2014). **i**. Biarc et al. (2004). **j**. Zeller et al. (2014). **k**. Weir et al. (2013). **l**. Sokol et al. (2008). **m**. Castellarin et al. (2012). **n**. Kostic et al. (2012). **o**. cf. Chen et al. (2012); Candela et al. (2014) (more abundant in CRC patients); Weir et al. (2013) (more abundant in healthy subjects).

| Phylum | Species | Abbrev. | Bioproject | Refs. |
| --- | --- | --- | --- | --- |
| *Model species for genera significantly higher in healthy patients<sup>a</sup>* |  |  |  |  |
| Bacteroidetes | *Bacteroides vulgatus* ATCC 8482 | Bvu | PRJNA13378 | c |
| Bacteroidetes | *Bacteroides uniformis* ATCC 8492 | Bun | PRJNA18195 | c |
| Firmicutes | *Roseburia intestinalis* L1-82 (DSM 14610) | Rin | PRJNA30005 | d |
| Bacteroidetes | *Alistipes indistinctus* YIT 12060 | Ain | PRJNA46373 | c |
| Firmicutes | *Eubacterium rectale* ATCC 33656 | Ere | PRJNA29071 | e |
| Proteobacteria | *Parasutterella excrementihominis* YIT 11859 | Pex | PRJNA48497 | f |
| *Model species for genera significantly higher in CRC patients<sup>a</sup>* |  |  |  |  |
| Bacteroidetes | *Porphyromonas gingivalis* W83 | Pgi | PRJNA48 | g |
| Proteobacteria | *Escherichia coli* NC101 | Eco | PRJNA47121 | c,h |
| Firmicutes | *Enterococcus faecalis* V583 | Efa | PRJNA57669 | c |
| Firmicutes | *Streptococcus infantarius* ATCC BAA-102 | Sin | PRJNA20527 | i |
| Firmicutes | *Peptostreptococcus stomatis* DSM 17678 | Pst | PRJNA34073 | j |
| Bacteroidetes | *Bacteroides fragilis* YCH46 | Bfr | PRJNA58195 | g |
| *Model species for protective co-abundance groups<sup>b</sup>* |  |  |  |  |
| Actinobacteria | *Bifidobacterium longum* NCC2705 | Blo | PRJNA57939 | g,k |
| Firmicutes | *Faecalibacterium prausnitzii* SL3/3 | Fpr | PRJNA39151 | e,l |
| *Model species for pro-carcinogenic co-abundance groups<sup>b</sup>* |  |  |  |  |
| Fusobacteria | *Fusobacterium nucleatum* ATCC 23726 | Fnu | PRJNA49043 | m,n |
| Bacteroidetes | *Prevotella copri* DSM 18205 | Pco | PRJNA30025 | k,o |
| Firmicutes | *Coprobacillus* sp. D7 | Csp | PRJNA32495 | h |

The datasets with a significant difference in 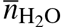 all show higher values for adenoma (5) or carcinoma (3) compared to normal tissue, up- expressed compared to down-expressed serum biomarker candidates (Jimenez et al., 2010), or secreted proteins detected in conditioned media (Yao et al., 2012). None of the datasets with a significant difference in 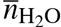 corresponds to a carcinoma / adenoma comparison.

Natural variability inherent in the heterogeneity of tumors, as well as differences in experimental design and technical analysis, may underlie the opposite trends in Z_C_ among some datasets that compare the same stages of cancer (e.g. carcinoma / adenoma). However, there is a preponderance of datasets with higher values of Z_C_ and 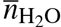 for the proteins with higher abundance in adenoma or carcinoma compared to normal tissue.

**Table III.** Species from a consensus microbial signature for CRC classification of fecal metagenomes (Zeller et al., 2014). Only species reported as having a log odds ratio larger than ±0.15 are listed here, together with strains and Bioproject IDs used as models in the present study.

| Species | Strain | Abbrv. | Bioproject |
| --- | --- | --- | --- |
| Higher in CRC patients |  |  |  |
| *Fusobacterium nucleatum* subsp. *vincentii* | ATCC 49256 | Fnv | PRJNA1419 |
| *Fusobacterium nucleatum* subsp. *animalis* | D11 | Fna | PRJNA32501 |
| *Peptostreptococcus stomatis* | DSM 17678 | Pst | PRJNA34073 |
| *Porphyromonas asaccharolytica* | DSM 20707 | Pas | PRJNA51745 |
| *Clostridium symbiosum* | ATCC 14940 | Csy | PRJNA18183 |
| *Clostridium hylemonae* | DSM 15053 | Chy | PRJNA30369 |
| *Lactobacillus salivarius* | ATCC 11741 | Lsa | PRJNA31503 |
| Higher in healthy patients |  |  |  |
| *Clostridium scindens* | ATCC 35704 | Csc | PRJNA18175 |
| *Eubacterium eligens* | ATCC 27750 | Eel | PRJNA29073 |
| *Methanospaera stadtmanae* | DSM 3091 | Mst | PRJNA15579 |
| *Phascolarctobacterium succinatutens* | YIT 12067 | Psu | PRJNA48505 |
| unclassified *Ruminococcus* sp. | ATCC 29149(*) | Rsp | PRJNA18179 |
| *Streptococcus salivarius* | SK126 | Ssa | PRJNA34091 |

\* *R. gnavus*

### 3.2 Compositional descriptions of microbial proteins

Summary data on microbial populations from four studies were selected for comparison here. First, in a study of 16S RNA of fecal microbiota, Wang et al. (2012) reported genera that are significantly increased or decreased in CRC compared to healthy patients. In order to compare the chemical compositions of the microbial population, single species with sequenced genomes were chosen to represent each of these genera (see Table 2). Where possible, the species selected are those mentioned by Wang et al. (2012) as being significantly altered, or are species reported in other studies to be present in healthy or cancer states (see Table 2).

In the second study considered (Zeller et al., 2014), changes in the metagenomic abundance of fecal microbiota associated with CRC were analyzed for their potential as a biosignature for cancer detection. The species shown in Fig. 1A of Zeller et al. (2014) with a log odds ratio greater than 0.15 were selected for comparison, and are listed in Table 3. Zeller et al. (2014) found a strong enrichment of *Fusobacterium* in cancer, consistent with previous reports (Kostic et al., 2012; Castellarin et al., 2012). In a third study, Candela et al. (2014) reported the findings of a network analysis that identified 5 microbial “co-abundance groups” at the genus level. As before, single representative species were selected in this study, and are listed in Table 2. Except for the presence of *Fusobacterium*, the co-abundance groups show little genus-level overlap with profiles derived from the previous two studies.

Finally, Table 4 lists the “best aligned strain” from Supplementary Dataset 5 of Feng et al. (2015) for all species shown there with negative enrichment in cancer, and for selected species with positive enrichment in cancer. Although every uniquely named strain given by Feng et al. (2015) was used in the comparisons here (n = 44; see Fig. 3D below), for clarity only the up-enriched species that appear in the calculated stability diagram (see Fig. 4D below) are listed in Table 4 and labeled in Fig. 3D. File `microbes.csv` in Dataset S1 contains the complete list of Bioproject IDs and calculated Z_C_ and 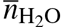 for all the microbial studies considered here.

**Table IV.** **Selected microbial species with negative or positive enrichment in cancer (Feng et al., 2015)**.

| Enriched Species | Abbrv. | Depleted Species | Abbrv. |
| --- | --- | --- | --- |
| *Bacteroides dorei* | Bdo | *Actinomyces viscosus* | Avi |
| *Bacteroides ovatus* | Bov | *Bifidobacterium animalis* | Ban |
| butyrate-producing bacterium SS3/4 | But | *Clostridium* sp. SS2/1 | Csp |
| *Clostridium asparagiforme* | Cas | *Ruminococcus* sp. 5_1_39BFAA | Rsp |
| *Fusobacterium* sp. oral taxon 370 | Fsp | *Streptococcus mutans* | Smu |
| *Lachnospiraceae* bacterium 3_1_57FAA_CT1 | L57 | *Streptococcus thermophilus* | Sth |
| *Paraprevotella clara* | Pcl |  |  |
| *Peptostreptococcus stomatis* | Pst |  |  |
| *Ruminococcaceae* bacterium D16 | Rba |  |  |

For each of the microbial species listed in Table 2–4, an overall protein composition was calculated by combining amino acid sequences of all proteins downloaded from the NCBI genome page associated with the Bioproject IDs shown in the Tables (see file `microbial.aa.csv` in the Dataset S1). This method does not account for actual protein abundance in organisms, and excludes any post-translational modifications. Calculation of the overall amino acid composition of proteins in this way is not an exact representation of the cellular protein composition, but provides a starting point for identifying environmental signals in protein composition. Mean amino acid composition, or amino acid frequencies deduced from microbial genomes, without weighting for actual protein abundance, has been used in many studies making evolutionary or environmental comparisons (Tekaia and Yeramian, 2006; Zeldovich et al., 2007; Brbić et al., 2015). More refined calculations of overall amino acid composition may be possible with genome-wide estimates of protein expression levels based on codon usage patterns (e.g. Moura et al., 2013; Brbić et al., 2015).

The water demand per residue 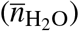 vs. oxidation state of carbon (Z_C_) in the overall amino acid compositions of proteins from all of the microbial species considered here are plotted in Fig. 1B and D, and for individual datasets in Fig. 3. As a group, the proteins in the microbes from cancer patients have somewhat lower Z_C_ than the healthy patients in the same study. The dataset from Feng et al. (2015) Fig. 3D shows a more complex distribution, where the microbes with a relative enrichment in healthy individuals form two clusters at high and low Z_C_. The *Fusobacterium* species identified in the studies of Zeller et al. (2014), Candela et al. (2014) and Feng et al. (2015) have the lowest Z_C_ of any microbial species considered here. The overall human protein composition is also plotted in Fig. 3, revealing a higher Z_C_ than any of the mean microbial proteins except for *Actinomyces viscosus* and *Bifidobacterium animalis*, identified in the study of Feng et al. (2015) (Fig. 3D). The tendency for microbial organisms to be composed of more reduced biomolecules than the host may reflect the relatively reducing conditions in the gut.

**Fig. 3.**
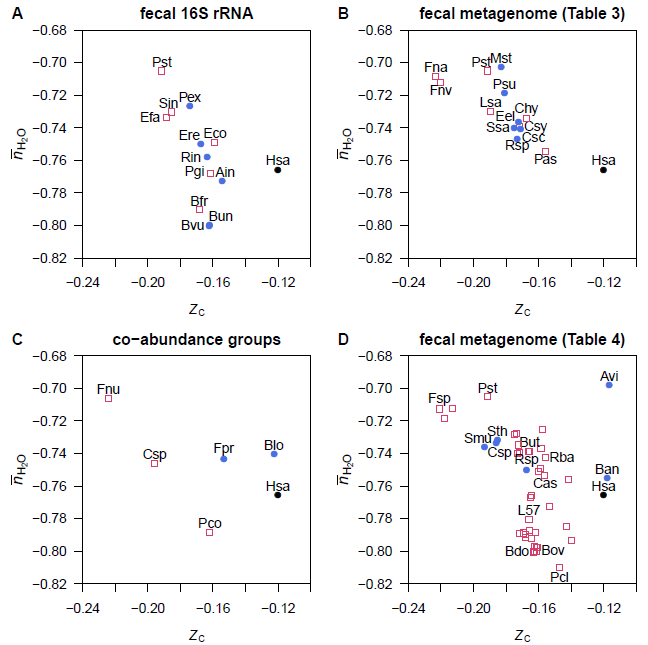
Average oxidation state of carbon (Z_C_) and water demand per residue 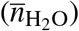 for overall amino acid compositions of proteins in genomes of normal-and cancer-associated microbes. Data are shown for representative species for (A) microbial genera identified in fecal 16s RNA (Wang et al., 2012; Table 2 top), (B) microbial signatures in fecal metagenomes (Zeller et al., 2014; Table 3), (C) microbial co-abundance groups (Candela et al., 2014; Table 2 bottom), and (D) best aligned strains to metagenomic linkage groups in fecal samples (Feng et al., 2015; Table 4). The location of the overall amino acid composition of proteins in humans (Hsa) is also shown.

### 3.3 Thermodynamic descriptions: background

Compositional comparisons by themselves do not yield physical models with relationships to biochemical conditions. A thermodynamic description can account for stoichiometric and energetic constraints and provide a richer interpretation of proteomic data in the context of tumor microenvironments.

By combining both stoichiometric and energetic variables, a thermodynamic description of proteomic data reveals possible biochemical constraints that may arise within cells and in tumor microenvironments. To give an example of how relative stabilities of up- and down-expressed proteins in a proteomic dataset can be calculated as a function of chemical potentials, consider Reaction R3 above written for the formation of one mole of MUC1. In order to compare proteins of different lengths, the formula of the protein is written per residue. The corresponding reaction is then

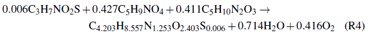

An expression for the chemical affinity (Kondepudi and Prigogine, 1998) of Reaction R4 is

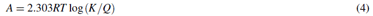

where the activity quotient *Q* is given by

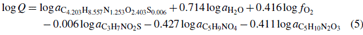

and the equilibrium constant is given by 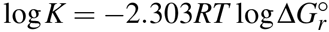, where 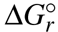 is the standard Gibbs energy of the reaction. As noted above, the standard Gibbs energies of species used to calculate 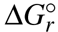 at *T* = 37°C are generated using amino acid group additivity for the proteins and published values for standard thermodynamic properties of the basis species in the reaction.

Here, the per-residue formulas of the proteins are given equal activities (1) and chemical activities of the amino acid basis species are set to nominal constant values (10^−4^), while 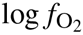 and 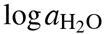 are used as exploratory variables. The ranges of these variables shown on the diagrams are selected in order to encompass the stability boundaries between groups of proteins differentially enriched in cancer and normal samples. There are combinations of chemical activities of basis species in Eq. (5) where the per-residue formation reaction have an equal affinity, indicating equal chemical stability of the proteins. Other combinations of chemical activities of basis species give the result that one protein-residue formula has a higher affinity than the others, indicating greater stability of this protein. This is the basis for the “maximum affinity method” for constructing stability diagrams described previously (Dick, 2008) and used below for microbial proteins.

### 3.4 Relative stability fields for microbial proteins

Stability diagrams are shown in Fig. 4A–D for the four sets of microbial proteins described above. The first diagram, representing significantly changed genera detected in fecal 16S rRNA (Wang et al., 2012; first part of Table 2, shows maximal stability fields for proteins from 5 species relatively enriched in healthy patients, and 3 species enriched in CRC patients. The other 4 proteins in the system are less stable than the others within the range of 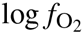 and 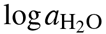 shown and do not appear on the diagram.

**Fig. 4.**
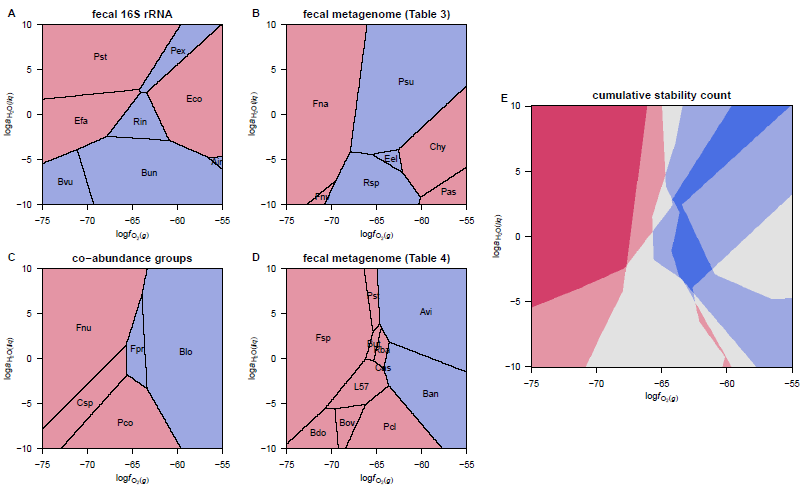
Maximal relative stability diagrams for overall microbial protein compositions. The diagrams show the range of oxygen fugacity and water activity (in log units: 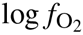 and 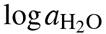) where the protein composition from the labeled microbial species has a higher affinity (lower Gibbs energy) of formation than the others. Blue and red shading designate microbes associated with normal and cancer samples, respectively. Plot (E) is a composite figure in which the intensity of shading corresponds to the number of overlapping cancer- or normal-enriched microbes in the preceding diagrams.

The relative positions of the stability fields in Fig. 4A are roughly aligned with the values of Z_C_ and 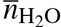 of the proteins; note for example the high-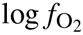 positions of the fields for the relatively high-Z_C_ *Escherichia coli* and *Alistipes indistinctus*, and the high-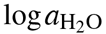 position of the field for the high-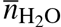 *Peptostreptococcus stomatis.* Except for *E. coli*, the proteins from the species associated with CRC in this dataset occupy the lower 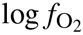 (reducing) and higher 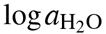 zones of this diagram.

In thermodynamic calculations for proteins from bacteria detected in fecal metagenomes (Zeller et al., 2014; Table 3), the mean protein compositions of 3 of 6 normal-enriched microbes and 4 of 7 cancer-enriched microbes exhibit maximal relative stability fields (Fig. 4B). Here, the cancer-associated proteins occupy the more reducing (*Fusobacterium nucleatum* subsp. vincentii and subsp. animalis) or more oxidizing (*Clostridium hylemonae, Porphyromonas asaccharolytica*) regions, while the proteins from bacteria more abundant in healthy individuals are relatively stable at moderate oxidation-reduction conditions.

For the bacterial species representing microbial co-abundance groups (Candela et al., 2014; second part of Table 2), all of the 5 mean protein compositions show up on the diagram. Here, the proteins from cancer-enriched bacteria are more stable at reducing conditions and those from normal-enriched microbes are stabilized by oxidizing conditions.

A stability diagram for proteins of bacteria identified in a second metagenomic study (Feng et al., 2015) shows a similar result (Fig. 3D) for the 11 overall protein compositions with highest stability at some point the diagram. These patterns in relative stability again reflect the differences in Z_C_ of the proteins, although in this case, a greater proportion of proteins (33 out of the 44 included in the calculations) are not found to have maximal stability fields. The resulting stability diagram is therefore a more limited portrayal of the available data.

Fig. 4E is a composite representation of the calculations, in which higher cumulative counts of maximal stability of proteins from bacteria enriched in normal and cancer samples in the four studies are represented by deeper blue and red shading, respectively. According to this diagram, the chemical conditions predicted to be most favorable for the formation of proteins in many bacteria enriched in CRC are characterized low 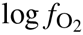. Proteins from bacteria that are abundant in healthy patients tend to be stabilized by moderate values of 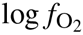. Despite the differences in experimental design and microbial identification between studies, the thermodynamic calculations reveal a shared pattern of relative stabilities among the four datasets considered here.

### 3.5 Relative stability fields for human proteins

Diagrams like those shown above that portray the maximally stable protein compositions are inadequate for analysis of larger datasets such as those generated in proteomic studies. It is apparent in Fig. 5 that only three different proteins up-expressed in cancer, from the 106 proteins in the `KWA+14` dataset (chromatin-binding proteins in carcinoma / adenoma), are maximally stable across a range of 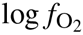. However, visual inspection reveals a differential sensitivity to oxygen fugacity in the whole dataset, with lower 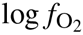 providing relatively higher potential for the formation of many of the up-expressed proteins in carcinoma samples. How can these responses be quantified in order to explore the data in multiple dimensions, including both 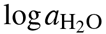 and 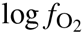?

In Fig. 5B, the difference in mean values of chemical affinity per residue of carcinoma and adenoma-associated proteins appears as a straight line as a function of 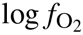. This linear behavior would translate to evenly-spaced iso-stability (as constant mean affinity difference) contours on a 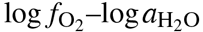 diagram. The weighted rank difference of affinities (see Methods), shown by the jagged curving line Fig. 5B, is a summary function that is more informative in changing chemical conditions. The variable slope is greatest near the zone of convergence for affinities of individual proteins, corresponding to the transition zone between groups of proteins. Two-dimensional iso-stability (as constant weighted rank difference of affinity) diagrams have curved and diversely spaced contours.

The diagrams in Fig. 6 portray weighted rank differences of chemical affinities of formation between groups of up- and down-expressed proteins reported for proteomic experiments. These combined depictions of stoichiometric and energetic differences constitute a theoretical prediction of the relative chemical (not conformational) stabilities of the proteins.

**Fig. 5.**
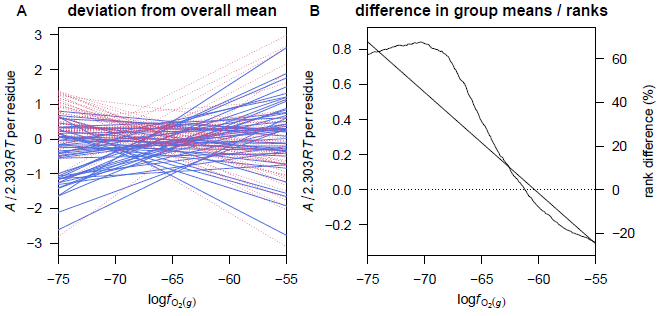
Calculated chemical affinities per residue of proteins in the KWA+14 dataset. Values for individual proteins as a function of 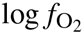 at 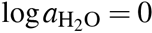 are shown in plot (A) as deviations from the mean value for all proteins. Healthy-and cancer-related proteins are indicated by solid blue and dashed red lines, respectively. Plot (B) shows the difference in mean value between healthy and cancer proteins (straight line and left-hand *y*-axis) and the weighted difference in sums of ranks of affinities as a percentage of maximum possible rank-sum difference (Eq. 3; jagged line and right-side *y*-axis). Positive values of affinity or rank-sum difference in plot (B) correspond to relatively greater stability of the cancer-related proteins.

The slopes of the equal-stability lines and the positions of the stability lines and the positions of the stability fields reflect the magnitude and sign of differences in Z_C_ and 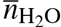. Figs. 6A–C show results for datasets that are dominated by differences in 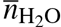; the nearly horizontal lines show that relative stabilities are accordingly more sensitive to 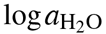. The second row depicts relative stabilities in the three datasets from Mikula et al. (2011), which have large changes in, sequentially, 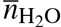, Z_C_, then both of these Table 1. Accordingly, the equal-stability lines for these datasets are closer to horizontal, closer to vertical, or have a more a diagonal trend (Fig. 6D–F).

The last row shows results for datasets that are characterized by large changes in Z_C_; the relative stabilities depend strongly on 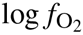. According to Fig. 6G, higher oxygen fugacity increases the relative potential for the formation of proteins up-expressed in cancer (dataset of Jankova et al. (2011)). However, using a dataset for up- and down-expressed chromatin-binding proteins in carcinoma (Knol et al., 2014), lower 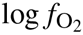 is predicted to promote formation of the proteins up-expressed in carcinoma. This is the opposite trend to that found for most of the other datasets with significant differences in Z_C_. These opposing trends might be attributed to different biochemical constraints on subcellular proteomes and whole-cell or whole-tissue proteomes during carcinogenesis. The full set of diagrams for all datasets listed in Table 1 is provided in Figure S1. It is notable that for the datasets where the relative stabilities are strongly a function of 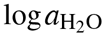 (sub-horizontal lines), the equal-stability lines are within a few log units of 0 (unit activity). Equal-stability lines that are diagonal often cross unit activity of H_2_O at a moderate value of 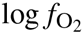, near −65 to −60 (see Figure S1). This could be indicative of a tendency for these proteomic transformations to be incompletely buffered by other redox reactions in the cell, and/or by liquid-like H_2_O with close to unit activity.

**Fig. 6.**
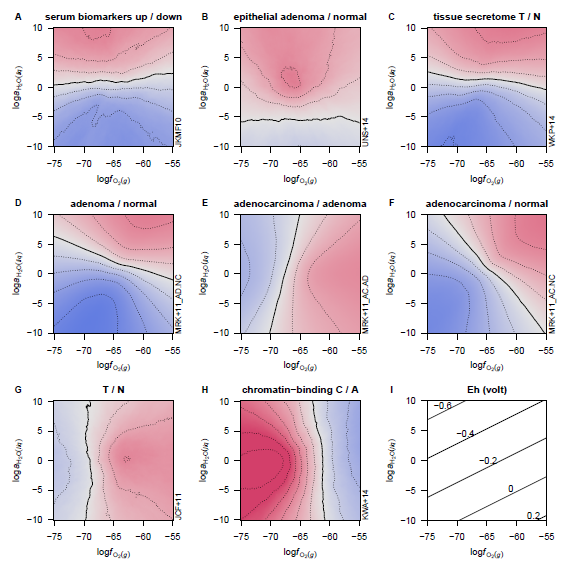
Weighted rank-sum comparisons of chemical affinities of formation of human proteins as a function of 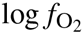 and 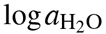. The solid lines indicate equal ranking of proteins in the “normal” and “cancer” groups (Table 1), and dotted contours are drawn at 10% increments of the maximum possible rank-sum difference. Blue and red areas correspond to higher ranking of cancer- and healthy-related proteins, respectively, with the intensity of the shading increasing up to 50% the maximum possible rank-sum difference. (For readers without a color copy: the stability fields for proteins up-expressed in cancer lie above (A-D), to the right of (E-G), or to the left of (H) the stability fields for proteins with higher expression in normal tissue.) Panel (I) shows calculated values of Eh over the same range of 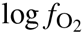 and 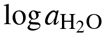 (cf. Reaction R4).

Effective values of oxidation-reduction potential (Eh) can be calculated by considering the water dissociation reaction, i.e.

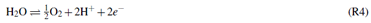

If one assumes that 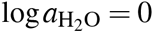 (unit water activity, as in an infinitely dilute solution), this reaction can be used to interconvert 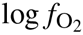, pH and pe (or, in conjunction with the Nernst equation, Eh) (e.g. Garrels and Christ, 1965, p. 176; Anderson, 2005, p. 363). However, in the approach proposed here for metastable equilibrium among proteins in a subcellular metabolic context, no such assumptions are made on the operational value of 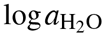, used as an internal indicator, not necessarily externally buffered by an aqueous solution. Consequently, the effective Eh is considered to be a function of variable 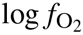 and 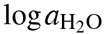, as shown in Fig. 6I for pH = 7.4 and *T* = 37°C. This comparison gives some perspective on operationally reasonable ranges of 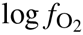 and 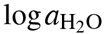.

The subcellular reduction potential monitored by the reduced glutathione (GSH) / oxidized glutathione disulfide (GSSG) couple ranges from ca. −260 mV for proliferating cells to ca. −170 mV for apoptotic cells (Schafer and Buettner, 2001), lying toward the middle part of the range of conditions shown in Fig. 6. A physiologically plausible Eh value of −0.2 V, corresponding to 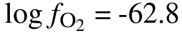 at unit activity of H_2_O, lies close to the stability transitions for many of the datasets considered here (see also Figure S1).

### 3.6 Comparison with inorganic basis species

Figures made using Basis I (inorganic basis species, e.g. Reaction R1) are provided in the Supplemental Information (human proteins: Figure S2; microbial proteins: Figure S3). The stability boundaries in 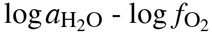 diagrams constructed using Basis I cluster around a common, positive slope, in contrast with the greater diversity of slopes appearing on the corresponding diagrams constructed using Basis II (Figure S1).

As noted above, all mathematically possible choices for the basis species of a system are thermodynamically valid, but it appears that Basis II affords a greater convenience for interpretation. That is, compared to to Basis I, Basis II yields a greater degree of separation of the effects of changing chemical potentials of H_2_O and O_2_ under the assumption that the activities of the remaining basis species (inorganic species in Basis I, or amino acids in Basis II) are held constant. However, it is also notable that two of the diagrams constructed using Basis I (Figure S2), unlike the others, have nearly horizontal equal-stability lines, showing that increasing activity of H_2_O at constant activity of CO_2_, NH_3_, H_2_S and fugacity of O_2_ gives an energetic advantage to the formation of potential up-expressed serum biomarkers (dataset `JKMF10`; Jimenez et al., 2010) and proteins up-expressed in an “epithelial cell signature” for adenoma (dataset `UNS+14`; Uzozie et al., 2014). These datasets are also included among those shown to have differential water demand using Basis II (Table 1; Figure S1). Based on the similar results for these datasets using different choices of chemical components, it can be suggested that the compositions of the differentially expressed proteins in these datasets are particularly indicative of changes in hydration potential.

## 4 DISCUSSION

Among 35 proteomic datasets considered here (Table 1), many have significantly higher values of average oxidation state of carbon (Z_C_) in proteins up-expressed in adenoma or carcinoma compared to normal tissue. While a decrease in oxidation state might be expected if biomacromolecular adaptation was driven by hypoxic conditions in tumors, the observed increase is more consistent with potentially oxidizing subcellular conditions that may accompany mitochondrial generation of ROS.

The datasets that show a negative change in Z_C_ (i.e. toward more reduced proteins) include one for the nuclear matrix fraction in chromosomal instability (CIN-type) CRC (Albrethsen et al., 2010), and one for the chromatin-binding fraction (Knol et al., 2014); both studies compared tissue samples between adenoma and carcinoma. Based on these results, it seems likely that particular subtypes of cancer and subfractions of cells have patterns of protein expression during carcinogenesis that are chemically distinct from the general trend toward proteins with higher oxidation state of carbon.

A couple of proteomic datasets are also available for stromal cells associated with tumor tissues. Data from one study (Mu et al., 2013) are consistent with the generally observed higher Z_C_ of protein in in tumors, but data from a pair of recent studies which analyzed cancer and stromal cells from the same set of tissues (Li et al., 2016; Peng et al., 2016) indicates that the proteins up-expressed in stromal cells, but not tumor cells, of adenoma are reduced compared to normal cells. Also, proteins up-expressed in tumor cells, but not stromal cells from carcinoma *in situ*, have a relatively oxidized composition (Table 1). If an opposing trend in Z_C_ between stromal and epithelial cells is indeed established, it might be evidence for a proteome-level response to metabolic coupling (Martinez-Outschoorn et al., 2014) between tissue compartments in cancer. The “lactate shuttle” that underlies this type of metabolic coupling can be characterized in part by the difference between oxidation state of lactate (Z_C_ = 0) and pyruvate (Z_C_ = 0.667) (Brooks, 2009). More work is needed to determine whether the fluxes of anabolic precursors and catabolic products between tissue compartments contribute to the differential oxidation states of carbon in proteins observed in cancer.

The datasets available for comparison of overall protein compositions of bacteria associated with normal and cancer states are characterized by lower Z_C_ in proteins of bacteria with higher abundance in cancer patients (Fig. 3), and consequently stabilization of proteins by lower oxygen fugacity (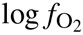; Fig. 4). This trend could be viewed as an adaptation of microbial communities to minimize the energetic costs of biomass synthesis in more reducing conditions. The opposite trends in Z_C_ for the human and bacterial proteins also raises the possibility that their mutual proteomic makeup is partially the result of a redox balance, or coupling.

Another major outcome of the compositional comparisons of human proteomes is the increase in water demand per residue 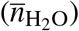 apparent in some datasets for CRC tissues and in a list of candidate biomarkers summarized in a literature review (Jimenez et al., 2010) (Table 1). Higher hydration levels in breast cancer tissues have been observed spectroscopically (Abramczyk et al., 2014), and it has been proposed that increased hydration plays a role in reversion to an embryological mode of growth (McIntyre, 2006). The thermodynamic calculations used to generate Fig. 6 support the possibility that higher water activity increases the potential for formation of the proteins up-expressed in cancer relative to normal tissue.

The conceptual basis for using 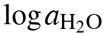 and 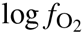 as indicators of the hydration and oxidation state of the system (Anderson, 2005) does not support a direct interpretation in terms of measurable concentrations. There are astronomical differences between theoretical values of oxygen fugacity and actual concentrations or partial pressures of oxygen (e.g. Anderson, 2005, p. 364-365). Partial pressures of oxygen in human arterial blood are around 90-100 mmHg, and approximate threshold values for physiological hypoxia include 10 mmHg for energy metabolism, 0.5 mmHg for mitochondrial oxidative phosphorylation, and 0.02 mmHg for full oxidation of cytochromes (Höckel and Vaupel, 2001). Assuming ideal mixing, the equivalent range of oxygen fugacities indicated by these measurements is 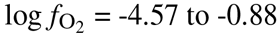, higher by far than the values that delimit the relative stabilities of cancer-and normal-enriched proteins computed here.

Likewise, the ranges of 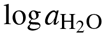 calculated here deviate tremendously from laboratory-based determination of water activity or hydration levels. Water activity in saturated protein solutions is not lower than 0.5 (Knezic et al., 2004), and recent experiments and extrapolations predict a range of ca. 0.600 to 0.650 for growth of various xerophilic and halophilic eukaryotes and prokaryotes (Stevenson et al., 2015). In general, cytoplasmic water activity is probably not greatly different from aqueous growth media, at 0.95 to 1 (Cayley et al., 2000). The theoretically computed transitions in relative stabilities between proteins from cancer and healthy tissues occur at much lower values of 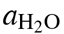 (ca. 10^−6^; Fig. 6B) or at values approaching 1, depending on the oxygen fugacity (Fig. 6; Figure S1).

Despite the difficulties in a quantitative interpretation, theoretical predictions of stabilization of cancer-related proteins by an increase in 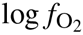 (e.g. Fig. 6D-G) can be interpreted qualitatively as corresponding with an increase in redox potential if log*a*_H_2_O_ is held constant (Fig. 6I). Alternatively, proteins up-expressed in cancer tissues in each of the datasets shown in Fig. 6A–G can be relatively stabilized along a trajectory of increasing both 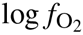 and 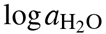 at constant redox potential near −0.2 V (Fig. 6I). Under this interpretation, global increases in both oxidation and hydration state are a general feature of the proteomic transformations in colorectal cancer.

## 5 CONCLUSION

An integrated picture of proteomic remodeling in cancer may benefit from accounting for the stoichiometric and energetic requirements of protein formation. This study has identified a strong shift toward higher average oxidation state of carbon in proteins that are more highly expressed in colorectal cancer. Importantly, this pattern is identified across multiple data sets, increasing confidence in its systematic nature. In some other data sets, a systematic change can be identified indicating greater water demand of human proteins in cancer compared to normal tissue.

The proteomic data can be theoretically linked to microenvironmental conditions using thermodynamic models, which give estimates of the oxidation- and hydration-potential limits for relative stability of groups of proteins. These calculations outline a path connecting the dynamic compositions of proteomes to biochemical measurements such as Eh, providing a new view of how proteomic transformations may be used as indicators of changing microenvironmental conditions. This approach can be used in conjunction with other datasets to characterize chemical changes in proteomes in different types of cancer and in the progression to metastasis.

## 6 ACKNOWLEDGEMENTS

Thanks to Greg Anderson for a helpful discussion about chemical components and Apar Prasad for giving many useful comments on the manuscript.

## REFERENCES

Abramczyk, H., Brozek-Pluska, B., Krzesniak, M., Kopec, M., and Morawiec-Sztandera, A. (2014). The cellular environment of cancerous human tissue. Interfacial and dangling water as a ‘hydration fingerprint”. Spectrochimica Acta, Part A: Molecular and Biomolecular Spectroscopy, 129(0):609–623. doi:10.1016/j.saa.2014.03.103.

Albarède, F. (2011). Oxygen fugacity. In Gargaud, M., Amils, R., Quintanilla, J. C., Cleaves, H. J. J., Irvine W. M., et al., editors, Encyclopedia of Astrobiology, pages 1196–1196. Springer, Berlin, Heidelberg. DOI:10.1007/978-3-642-11274-4_4021.

Alberty, R. A. (2004). Equilibrium concentrations for pyruvate dehydrogenase and the citric acid cycle at specified concentrations of certain coenzymes. Biophysical Chemistry, 109(109):73–84. DOI:10.1016/j.bpc.2003.10.019.

Albrethsen, J., Knol, J. C., Piersma, S. R., Pham, T. V., de Wit, M., et al. (2010). Subnuclear proteomics in colorectal cancer: Identification of proteins enriched in the nuclear matrix fraction and regulation in adenoma to carcinoma progression. Molecular & Cellular Proteomics, 9(9):988–1005. DOI:10.1074/mcp.M900546-MCP200.

Amend, J. P., LaRowe, D. E., McCollom, T. M., and Shock, E. L. (2013). The energetics of organic synthesis inside and outside the cell. Philosophical Transactions of the Royal Society, B: Biological Sciences, 368(368):2012–0255. DOI:10.1098/rstb.2012.0255.

Amend, J. P. and Shock, E. L. (1998). Energetics of amino acid synthesis in hydrothermal ecosystems. Science, 281(281):1659–1662. DOI:10.1126/science.281.5383.1659.

Anderson, G. M. (2005). Thermodynamics of Natural Systems. Cambridge University Press, Cambridge, 2nd edition. URL http://www.worldcat.org/oclc/474880901

Arndt, S., JØrgensen, B. B., LaRowe, D. E., Middelburg, J. J., Pancost, R. D., et al. (2013). Quantifying the degradation of organic matter in marine sediments: A review and synthesis. Earth-Science Reviews, 123(123):53–86. DOI:10.1016/j.earscirev.2013.02.008.

Besson, D., Pavageau, A.-H., Valo, I., Bourreau, A., Belanger, A., et al. (2011). A quantitative proteomic approach of the different stages of colorectal cancer establishes OLFM4 as a new nonmetastatic tumor marker. Molecular & Cellular Proteomics, 10(10):M111.009712. DOI:10.1074/mcp.M111.009712.

Bethke, C. M. (2008). Geochemical and Biogeochemical Reaction Modeling. Cambridge University Press, 2nd edition.

Biarc, J., Nguyen, I. S., Pini, A., Gossè, F., Richert, S., et al. (2004). Carcinogenic properties of proteins with pro-inflammatory activity from Streptococcus infantarius (formerly S.bovis). Carcinogenesis, 25(8):1477–1484. DOI:10.1093/carcin/bgh091.

Bohutskyi, P., Chow, S., Ketter, B., Betenbaugh, M. J., and Bouwer, E. J. (2015). Prospects for methane production and nutrient recycling from lipid extracted residues and whole Nannochloropsis salina using anaerobic digestion. Applied Energy, 154:718–731. DOI:10.1016/j.apenergy.2015.05.069.

Borak, B., Ort, D. R., and Burbaum, J. J. (2013). Energy and carbon accounting to compare bioenergy crops. Current Opinion in Biotechnology, 24(24):369–375. DOI:10.1016/j.copbio.2013.02.018.

Brbić, M., Warnecke, T., Kriško, A., and Supek, F. (2015). Global shifts in genome and proteome composition are very tightly coupled. Genome Biology and Evolution, 7(7):1519–1532. DOI:10.1093/gbe/evv088.

Brooks, G. A. (2009). Cell-cell and intracellular lactate shuttles. Journal of Physiology, 587(23):5591–5600. DOI:10.1113/jphysiol.2009.178350.

Callen, H. B. (1985). Thermodynamics and an Introduction to Thermostatistics. John Wiley & Sons, 2nd edition. URL http://www.worldcat.org/oclc/11916089

Candela, M., Turroni, S., Biagi, E., Carbonero, F., Rampelli, S., et al. (2014). Inflammation and colorectal cancer, when microbiota-host mutualism breaks. World Journal of Gastroenterology, 20(4):908–922. DOI:10.3748/wjg.v20.i4.908.

Castellarin, M., Warren, R. L., Freeman, J. D., Dreolini, L., Krzywinski, M., et al. (2012). Fusobacterium nucleatum infection is prevalent in human colorectal carcinoma. Genome Research, 22(2):299–306. DOI:10.1101/gr.126516.111.

Cayley, D. S., Guttman, H. J., and Record Jr, M. T. (2000). Biophysical characterization of changes in amounts and activity of Escherichia coli cell and compartment water and turgor pressure in response to osmotic stress. Biophysical Journal, 78(4):1748–1764. DOI:10.1016/S0006-0006(00)6726-76726.

Chen, W., Liu, F., Ling, Z., Tong, X., and Xiang, C. (2012). Human intestinal lumen and mucosa-associated microbiota in patients with colorectal cancer. PLoS ONE, 7(7):e39743. DOI:10.1371/journal.pone.0039743.

Damer, B. and Deamer, D. (2015). Coupled phases and combinatorial selection in fluctuating hydrothermal pools: A scenario to guide experimental approaches to the origin of cellular life. Life, 5(5):872–887. DOI:10.3390/life5010872.

Davies, P. C. W., Demetrius, L., and Tuszynski, J. A. (2011). Cancer as a dynamical phase transition. Theoretical Biology and Medical Modelling, 8:30. DOI:10.1186/1742-4682-8-30.

de Wit, M., Fijneman, R. J. A., Verheul, H. M. W., Meijer, G. A., and Jimenez, C. R. (2013). Proteomics in colorectal cancer translational research: Biomarker discovery for clinical applications. Clinical Biochemistry, 46(46):466–479. DOI:10.1016/j.clinbiochem.2012.10.039.

de Wit, M., Kant, H., Piersma, S. R., Pham, T. V., Mongera, S., et al. (2014). Colorectal cancer candidate biomarkers identified by tissue secretome proteome profiling. Journal of Proteomics, 99:26–39. DOI:10.1016/j.jprot.2014.01.001.

DeBerardinis, R. J. and Cheng, T. (2010). Q’s next: The diverse functions of glutamine in metabolism, cell biology and cancer. Oncogene, 29(3):313–324. DOI:10.1038/onc.2009.358.

Dick, J. M. (2008). Calculation of the relative metastabilities of proteins using the CHNOSZ software package. Geochemical Transactions, 9:10. DOI:10.1186/1467-4866-9-10.

Dick, J. M. (2014). Average oxidation state of carbon in proteins. Journal of the Royal Society Interface, 11:2013–1095. DOI:10.1098/rsif.2013.1095.

Dick, J. M., LaRowe, D. E., and Helgeson, H. C. (2006). Temperature, pressure, and electrochemical constraints on protein speciation: Group additivity calculation of the standard molal thermodynamic properties of ionized unfolded proteins. Biogeosciences, 3(3):311–336. DOI:10.5194/bg-3-311-2006.

Duncan, S. H., Hold, G. L., Barcenilla, A., Stewart, C. S., and Flint, H. J. (2002). Roseburia intestinalis sp. nov., a novel saccharolytic, butyrate-producing bacterium from human faeces. International Journal of Systematic and Evolutionary Microbiology, 52(5):1615–1620. DOI:10.1099/ijs.0.02143-0.

Enver, T., Pera, M., Peterson, C., and Andrews, P. W. (2009). Stem cell states, fates, and the rules of attraction. Cell Stem Cell, 4(4):387–397. DOI:10.1016/j.stem.2009.04.011.

Fekete, J., Sajgó, C., Kramarics, A., Eke, Z., Kovács, K., et al. (2012). Aquathermolysis of humic and fulvic acids: Simulation of organic matter maturation in hot thermal waters. Organic Geochemistry, 53:109–118. DOI:10.1016/j.orggeochem.2012.07.005.

Feng, Q., Liang, S., Jia, H., Stadlmayr, A., Tang, L., et al. (2015). Gut microbiome development along the colorectal adenoma-carcinoma sequence. Nature Communications, 6:6528. DOI:10.1038/ncomms7528.

Garrels, R. M. and Christ, C. L. (1965). Solutions, Minerals, and Equilibria. Harper & Row, New York. URL http://www.worldcat.org/oclc/517586

Gibbs, J. W. (1875). On the equilibrium of heterogeneous substances (first part). Transactions of the Connecticut Academy of Arts and Sciences, 3:108–248. URL http://archive.org/details/Onequilibriumhe00Gibb.

Grzegorczyk, T. M., Meaney, P. M., Kaufman, P. A., di Florio-Alexander, R. M., and Paulsen, K. D. (2012). Fast 3-D tomographic microwave imaging for breast cancer detection. IEEE Transactions on Medical Imaging, 31(8):1584–1592. DOI:10.1109/TMI.2012.2197218.

Gupta, V., Ganegoda, H., Engelhard, M. H., Terry, J., and Linford, M. R. (2014). Assigning oxidation states to organic compounds via predictions from X-ray photoelectron spectroscopy: A discussion of approaches and recommended improvements. Journal of Chemical Education, 91(91):232–238. DOI:10.1021/ed400401c.

Guzy, R. D. and Schumacker, P. T. (2006). Oxygen sensing by mitochondria at complex III: The paradox of increased reactive oxygen species during hypoxia. Experimental Physiology, 91(91):807–819. DOI:10.1113/expphysiol.2006.033506.

Halkides, C. J. (2000). Assigning and using oxidation numbers in biochemistry lecture courses. Journal of Chemical Education, 77(77):1428–1432. DOI:10.1021/ed077p1428.

Hansen, L. D., Hopkin, M. S., Rank, D. R., Anekonda, T. S., Breidenbach, R. W., et al. (1994). The relation between plant growth and respiration: A thermodynamic model. Planta, 194(194):77–85. DOI:10.1007/BF00201037.

Häussinger, D. (1996). The role of cellular hydration in the regulation of cell function. Biochemical Journal, 313:697–710. DOI:10.1042/bj3130697.

Helgeson, H. C. (1970). Description and interpretation of phase relations in geochemical processes involving aqueous solutions. American Journal of Science, 268(5):415–438. DOI:10.2475/ajs.268.5.415.

Helgeson, H. C., Kirkham, D. H., and Flowers, G. C. (1981). Theoretical prediction of the thermodynamic behavior of aqueous electrolytes at high pressures and temperatures: IV. Calculation of activity coefficients, osmotic coefficients, and apparent molal and standard and relative partial molal properties to 600°C and 5 Kb. American Journal of Science, 281(10):1249–1516. DOI:10.2475/ajs.281.10.1249.

Helgeson, H. C., Richard, L., McKenzie, W. F., Norton, D. L., and Schmitt, A. (2009). A chemical and thermodynamic model of oil generation in hydrocarbon source rocks. Geochimica et Cosmochimica Acta, 73(73):594–695. DOI:10.1016/j.gca.2008.03.004.

Hendrickson, J. B., Cram, D. J., and Hammond, G. S. (1970). Organic Chemistry. McGraw-Hill, New York, 3rd edition. URL www.worldcat.org/oclc/78308

Hiller, K. and Metallo, C. M. (2013). Profiling metabolic networks to study cancer metabolism. Current Opinion in Biotechnology, 24(24):60–68. DOI:10.1016/j.copbio.2012.11.001.

Höckel, M. and Vaupel, P. (2001). Tumor hypoxia: Definitions and current clinical, biologic, and molecular aspects. Journal of the National Cancer Institute, 93(93):266–276. DOI:10.1093/jnci/93.4.266.

Hutter, D. E., Till, B. G., and Greene, J. J. (1997). Redox state changes in density-dependent regulation of proliferation. Experimental Cell Research, 232(2):435–438. DOI:10.1006/excr.1997.3527.

Ihaka, R., Murrell, P., Hornik, K., Fisher, J. C., and Zeileis, A. (2015). colorspace: Color Space Manipulation. R package version 1.2–6.

Jankova, L., Chan, C., Fung, C. L. S., Song, X., Kwun, S. Y., et al. (2011). Proteomic comparison of colorectal tumours and non-neoplastic mucosa from paired patient samples using iTRAQ mass spectrometry. Molecular Biosystems, 7:2997–3005. DOI:10.1039/C1MB05236E.

Jimenez, C. R., Knol, J. C., Meijer, G. A., and Fijneman, R. J. A. (2010). Proteomics of colorectal cancer: Overview of discovery studies and identification of commonly identified cancer-associated proteins and candidate CRC serum markers. Journal of Proteomics, 73(10):1873–1895. DOI:10.1016/j.jprot.2010.06.004.

Johnson, J. W., Oelkers, E. H., and Helgeson, H. C. (1992). SUPCRT92: A software package for calculating the standard molal thermodynamic properties of minerals, gases, aqueous species, and reactions from 1 to 5000 bar and 0 to 1000°C. Computers & Geosciences, 18(7):899–947. DOI:10.1016/0098-3004(92)90029-Q.

Kang, U.-B., Yeom, J., Kim, H.-J., Kim, H., and Lee, C. (2012). Expression profiling of more than 3500 proteins of MSS-type colorectal cancer by stable isotope labeling and mass spectrometry. Journal of Proteomics, 75(75):3050–3062. DOI:10.1016/j.jprot.2011.11.021.

Kelley, K. K. (1960). Contributions to the Data in Theoretical Metallurgy XIII: High Temperature Heat Content, Heat Capacities and Entropy Data for the Elements and Inorganic Compounds. Bulletin 584. U. S. Bureau of Mines. URL http://www.worldcat.org/oclc/693388901.

Kim, H.-J., Kang, U.-B., Lee, H., Jung, J.-H., Lee, S.-T., et al. (2012). Profiling of differentially expressed proteins in stage IV colorectal cancers with good and poor outcomes. Journal of Proteomics, 75(75):2983–2997. DOI:10.1016/j.jprot.2011.12.002.

Kinzler, K. W. and Vogelstein, B. (1996). Lessons from hereditary colorectal cancer. Cell, 87(2):159–170. DOI:10.1016/S0092-8674(00)81333-1.

Knezic, D., Zaccaro, J., and Myerson, A. S. (2004). Thermodynamic properties of supersaturated protein solutions. Crystal Growth & Design, 4(1):199–208. DOI:10.1021/cg034072o.

Knol, J. C., de Wit, M., Albrethsen, J., Piersma, S. R., Pham, T. V., et al. (2014). Proteomics of differential extraction fractions enriched for chromatin-binding proteins from colon adenoma and carcinoma tissues. Biochimica et Biophysica Acta (BBA)-Proteins and Proteomics, 1844(5):1034–1043. DOI:10.1016/j.bbapap.2013.12.006.

Kondepudi, D. K. and Prigogine, I. (1998). Modern Thermodynamics: From Heat Engines to Dissipative Structures. John Wiley & Sons, New York. URL http://www.worldcat.org/oclc/38055900.

Kostic, A. D., Gevers, D., Pedamallu, C. S., Michaud, M., Duke, F., et al. (2012). Genomic analysis identifies association of Fusobacterium with colorectal carcinoma. Genome Research, 22(2):292–298. DOI:10.1101/gr.126573.111.

Kroll, J. H., Donahue, N. M., Jimenez, J. L., Kessler, S. H., Canagaratna, M. R., et al. (2011). Carbon oxidation state as a metric for describing the chemistry of atmospheric organic aerosol. Nature Chemistry, 3(3):133–139. DOI:10.1038/NCHEM.948.

LaRowe, D. E. and Dick, J. M. (2012). Calculation of the standard molal thermodynamic properties of crystalline peptides. Geochimica et Cosmochimica Acta, 80:70–91. DOI:10.1016/j.gca.2011.11.041.

Lazaridis, T. and Karplus, M. (2003). Thermodynamics of protein folding: A microscopic view. Biophysical Chemistry, 100(1-3):367–395. DOI:10.1016/S0301-4622(02)00293-4.

Levy, Y. and Onuchic, J. N. (2006). Water mediation in protein folding and molecular recognition. Annual Review of Biophysics and Biomolecular Structure, 35:389–415. DOI:10.1146/annurev.biophys.35.040405.102134.

Li, M., Peng, F., Li, G., Fu, Y., Huang, Y., et al. (2016). Proteomic analysis of stromal proteins in different stages of colorectal cancer establishes Tenascin-C as a stromal biomarker for colorectal cancer metastasis. Oncotarget, 5(5). DOI:10.18632/oncotarget.9362.

Loock, H.-P. (2011). Expanded definition of the oxidation state. Journal of Chemical Education, 88(88):283–282. DOI:10.1021/ed1005213.

Louis, P. and Flint, H. J. (2007). Development of a semiquantitative degenerate real-time PCR-based assay for estimation of numbers of butyryl-coenzyme A (CoA) CoA transferase genes in complex bacterial samples. Applied and Environmental Microbiology, 73(6):2009–2012. DOI:10.1128/AEM.02561-06.

Martinez-Outschoorn, U. E., Lisanti, M. P., and Sotgia, F. (2014). Catabolic cancer-associated fibroblasts transfer energy and biomass to anabolic cancer cells, fueling tumor growth. Seminars in Cancer Biology, 25:47–60. DOI:10.1016/j.semcancer.2014.01.005.

Martínez-Aguilar, J., Chik, J., Nicholson, J., Semaan, C., McKay, M. J., et al. (2013). Quantitative mass spectrometry for colorectal cancer proteomics. Proteomics: Clinical Applications, 7(1-2):42–54. DOI:10.1002/prca.201200080.

Masiello, C. A., Gallagher, M. E., Randerson, J. T., Deco, R. M., and Chadwick, O. A. (2008). Evaluating two experimental approaches for measuring ecosystem carbon oxidation state and oxidative ratio. Journal of Geophysical Research, 113(G3):G03010. DOI:10.1029/2007JG000534.

May, P. M., and Murray, K. (2001). Database of chemical reactions designed to achieve thermodynamic consistency automatically. Journal of Chemical & Engineering Data, 46(5):1035–1040x. DOI:10.1021/je000246j.

McIntyre, G. I. (2006). Cell hydration as the primary factor in carcinogenesis: A unifying concept. Medical Hypotheses, 66(3):518–526. DOI:10.1016/j.mehy.2005.09.022.

Mikula, M., Rubel, T., Karczmarski, J., Goryca, K., Dadlez, M., et al. (2011). Integrating proteomic and transcriptomic high-throughput surveys for search of new biomarkers of colon tumors. Functional and Integrative Genomics, 11(11):215–224. DOI:10.1007/s10142-010-0200-5.

Morel, F. M. M. and Hering, J. G. (1993). Principles and Applications of Aquatic Chemistry. John Wiley & Sons.

Moura, A., Savageau, M. A., and Alves, R. (2013). Relative amino acid composition signatures of organisms and environments. PLoS ONE, 8(8):e77319. DOI:10.1371/journal.pone.0077319.

Mu, Y., Chen, Y., Zhang, G., Zhan, X., Li, Y., et al. (2013). Identification of stromal differentially expressed proteins in the colon carcinoma by quantitative proteomics. Electrophoresis, 34(34):1679–1692. DOI:10.1002/elps.201200596.

Murphy, M. P. (2009). How mitochondria produce reactive oxygen species. Biochemical Journal, 417:1–13. DOI:10.1042/bj20081386.

Nagai, F., Morotomi, M., Sakon, H., and Tanaka, R. (2009). Parasutterella excrementihominis gen. nov., sp. nov., a member of the family Alcaligenaceae isolated from human faeces. International Journal of Systematic and Evolutionary Microbiology 59(59):1793–1797. DOI:10.1099/ijs.0.002519-0.

Pace, N. R. (1991). Origin of life: Facing up to the physical setting. Cell, 65(65):531–533. DOI:10.1016/0092-8674(91)90082-A.

Peng, F., Huang, Y., Li, M.-Y., Li, G.-Q., Huang, H.-C., et al. (2016). Dissecting characteristics and dynamics of differentially expressed proteins during multistage carcinogenesis of human colorectal cancer. World Journal of Gastroenterology, 22(22):4515–4528. DOI:10.3748/wjg.v22.i18.4515.

R Core Team (2016). R: A Language and Environment for Statistical Computing. R Foundation for Statistical Computing, Vienna, Austria.

Ravi Kanth, M. V. S. R., Pushpavanam, S., Narasimhan, S., and Narasimha, M. B. (2014). A robust and efficient algorithm for computing reactive equilibria in single and multiphase systems. Industrial & Engineering Chemistry Research, 53(53):15278–15286. DOI:10.1021/ie502639a.

Riedel, T., Biester, H., and Dittmar, T. (2012). Molecular fractionation of dissolved organic matter with metal salts. Environmental Science & Technology, 46(46):4419–4426. DOI:10.1021/es203901u.

Rietman, E. A., Platig, J., Tuszynski, J. A., and Lakka Klement, G. (2016). Thermodynamic measures of cancer: Gibbs free energy and entropy of protein-protein interactions. Journal of Biological Physics, pages 1–12. DOI:10.1007/s10867-016-9410-y.

Rogmann, J. J. (2013). orddom: Ordinal Dominance Statistics. R package version 3.1.

Russell, M. J. and Hall, A. J. (1997). The emergence of life from iron monosulphide bubbles at a submarine hydrothermal redox and pH front. Journal of the Geological Society, 154:377–402. DOI:10.1144/gsjgs.154.3.0377.

Schafer, F. Q. and Buettner, G. R. (2001). Redox environment of the cell as viewed through the redox state of the glutathione disulfide/glutathione couple. Free Radical Biology and Medicine, 30(11):1191–1212. DOI:10.1016/S0891-5849(01)00480-4.

Schedin, P. and Elias, A. (2004). Multistep tumorigenesis and the microenvironment. Breast Cancer Research, 6(2):93–101. DOI:10.1186/bcr772.

Semenza, G. L. (2008). Tumor metabolism: Cancer cells give and take lactate. Journal of Clinical Investigation, 118(12):3835–3837. DOI:10.1172/JCI37373.

Sethi, M. K., Thaysen-Andersen, M., Kim, H., Park, C. K., Baker, M. S., et al. (2015). Quantitative proteomic analysis of paired colorectal cancer and non-tumorigenic tissues reveals signature proteins and perturbed pathways involved in CRC progression and metastasis. Journal of Proteomics, 126:54–67. DOI:10.1016/j.jprot.2015.05.037.

Shock, E. and Canovas, P. (2010). The potential for abiotic organic synthesis and biosynthesis at seafloor hydrothermal systems. Geofluids, 10(1-2):161–192. DOI:10.1111/j.1468-8123.2010.00277.x.

Sokol, H., Pigneur, B., Watterlot, L., Lakhdari, O., Bermiidez-Humaran, L. G., et al. (2008). Faecalibacterium prausnitzii is an anti-inflammatory commensal bacterium identified by gut microbiota analysis of Crohn disease patients. Proceedings of the National Academy of Sciences of the United States of America, 105(43):16731–16736. DOI:10.1073/pnas.0804812105.

Stevenson, A., Cray, J. A., Williams, J. P., Santos, R., Sahay, R., et al. (2015). Is there a common water-activity limit for the three domains of life? ISME Journal, 9(6):1333–1351. DOI:10.1038/ismej.2014.219.

Tanger, J. C. IV. and Helgeson, H. C. (1988). Calculation of the thermodynamic and transport properties of aqueous species at high pressures and temperatures: Revised equations of state for the standard partial molal properties of ions and electrolytes. American Journal of Science, 288(1):19–98. DOI:10.2475/ajs.288.1.19.

Tekaia, F. and Yeramian, E. (2006). Evolution of proteomes: Fundamental signatures and global trends in amino acid compositions. BMC Genomics, 7:307. DOI:10.1186/1471-2164-7-307.

Toole, B. P. (2002). Hyaluronan promotes the malignant phenotype. Glycobiology, 12(12):37R–42R. DOI:10.1093/glycob/12.3.37R.

Uzozie, A., Nanni, P., Staiano, T., Grossmann, J., Barkow-Oesterreicher, S., et al. (2014). Sorbitol dehydrogenase overexpression and other aspects of dysregulated protein expression in human precancerous colorectal neoplasms: A quantitative proteomics study. Molecular & Cellular Proteomics, 13(5):1198–1218. DOI:10.1074/mcp.M113.035105.

VanBriesen, J. M. and Rittmann, B. E. (1999). Modeling speciation effects on biodegradation in mixed metal/chelate systems. Biodegradation, 10(10):315–330. DOI:10.1023/A:1008375722721.

Wagman, D. D., Evans, W. H., Parker, V. B., Schumm, R. H., Halow, I., et al. (1982). The NBS tables of chemical thermodynamic properties. Selected values for inorganic and C_1_ and C_2_ organic substances in SI units. Journal of Physical and Chemical Reference Data, 11(11):1–392. DOI:10.1063/1.555661.

Wang, T., Cai, G., Qiu, Y., Fei, N., Zhang, M., et al. (2012). Structural segregation of gut microbiota between colorectal cancer patients and healthy volunteers. ISME Journal, 6(2):320–329. DOI:10.1038/ismej.2011.109.

Watanabe, M., Takemasa, I., Kawaguchi, N., Miyake, M., Nishimura, N., et al. (2008). An application of the 2-nitrobenzenesulfenyl method to proteomic profiling of human colorectal carcinoma: A novel approach for biomarker discovery. Proteomics: Clinical Applications, 2(2):925–935. DOI:10.1002/prca.200780111.

Wedberg, R., Abildskov, J., and Peters, G. H. (2012). Protein dynamics in organic media at varying water activity studied by molecular dynamics simulation. Journal of Physical Chemistry B, 116(116):2575–2585. DOI:10.1021/jp211054u.

Weir, T. L., Manter, D. K., Sheflin, A. M., Barnett, B. A., Heuberger, A. L., et al. (2013). Stool microbiome and metabolome differences between colorectal cancer patients and healthy adults. PLoS ONE, 8(8):e70803. DOI:10.1371/journal.pone.0070803.

Wiśniewski, J. R., Duś-Szachniewicz, K., Ostasiewicz, P., Zióikowski, P., Rakus, D., et al. (2015). Absolute proteome analysis of colorectal mucosa, adenoma, and cancer reveals drastic changes in fatty acid metabolism and plasma membrane transporters. Journal of Proteome Research, 14(14):4005–4018. DOI:10.1021/acs.jproteome.5b00523.

Wiśniewski, J. R., Ostasiewicz, P., Duś, K., ZieliĔska, D. F., Gnad, F., et al. (2012). Extensive quantitative remodeling of the proteome between normal colon tissue and adenocarcinoma. Molecular Systems Biology, 8(8):611. DOI:10.1038/msb.2012.44.

Xie, L.-Q., Zhao, C., Cai, S.-J., Xu,Y., Huang, L.-Y., et al. (2010). Novel proteomic strategy reveal combined α 1 antitrypsin and cathepsin D as biomarkers for colorectal cancer early screening. Journal of Proteome Research, 9(9):4701–4709. DOI:10.1021/pr100406z.

Yang, H.-Y., Chay, K.-O., Kwon, J., Kwon, S.-O., Park, Y.-K., et al. (2013). Comparative proteomic analysis of cysteine oxidation in colorectal cancer patients. Molecules and Cells, 35(35):533–542. DOI:10.1007/s10059-013-0058-1.

Yao, L., Lao, W., Zhang, Y., Tang, X., Hu, X., et al. (2012). Identification of EFEMP2 as a serum biomarker for the early detection of colorectal cancer with lectin affinity capture assisted secretome analysis of cultured fresh tissues. Journal of Proteome Research, 11(11):3281–3294. DOI:10.1021/pr300020p.

Yeh, C.-C., Lai, C.-Y., Hsieh, L.-L., Tang, R., Wu, F.-Y., et al. (2010). Protein carbonyl levels, glutathione S-transferase polymorphisms and risk of colorectal cancer. Carcinogenesis, 31(31):228–233. DOI:10.1093/carcin/bgp286.

Zeileis, A., Hornik, K., and Murrell, P. (2009). Escaping RGBland: Selecting colors for statistical graphics. Computational Statistics and Data Analysis, 53(53):3259–3270. DOI:10.1016/j.csda.2008.11.033.

Zeldovich, K. B., Berezovsky, I. N., and Shakhnovich, E. I. (2007). Protein and DNA sequence determinants of thermophilic adaptation. PLoS Computational Biology, 3(3):62–72. DOI:10.1371/journal.pcbi.0030005.

Zeller, G., Tap, J., Voigt, A. Y., Sunagawa, S., Kultima, J. R., et al. (2014). Potential of fecal microbiota for early-stage detection of colorectal cancer. Molecular Systems Biology, 10(11):766. DOI:10.15252/msb.20145645.

Zhang, Y., Ye, Y., Shen, D., Jiang, K., Zhang, H., et al. (2010). Identification of transgelin-2 as a biomarker of colorectal cancer by laser capture microdissection and quantitative proteome analysis. Cancer Science, 101(101):523–529. DOI:10.1111/j.1349-7006.2009.01424.x.

